# Modulation of root growth by nutrient-defined fine-tuning of polar auxin transport

**DOI:** 10.1101/2020.06.19.160994

**Authors:** Krisztina Ötvös, Marco Marconi, Andrea Vega, Jose O’ Brien, Alexander Johnson, Rashed Abualia, Livio Antonielli, Juan Carlos Montesinos, Yuzhou Zhang, Shutang Tan, Candela Cuesta, Christina Artner, Eleonore Bouguyon, Alain Gojon, Jirí Friml, Rodrigo A. Gutiérrez, Krzysztof Wabnik, Eva Benková

## Abstract

Nitrogen is an essential macronutrient and its availability in soil plays a critical role in plant growth, development and impacts agricultural productivity. Plants have evolved different strategies to sense and respond to heterogeneous nitrogen distribution. Modulating root system architecture, including primary root growth and branching, is among the most essential plant adaptions to ensure adequate nitrogen acquisition. However, the immediate molecular pathways coordinating the adjustment of root growth in response to varying nitrogen sources are poorly understood. Here, using a combination of physiological, live *in vivo* high- and super resolution imaging, we describe a novel adaptation strategy of root growth on available nitrogen source. We show that growth, *i.e*. tissue-specific cell division and elongation rates are fine-tuned by modulating auxin flux within and between tissues. Changes in auxin redistribution are achieved by nitrogen source dependent post-translational modification of PIN2, a major auxin efflux carrier, at an uncharacterized, evolutionary conserved phosphosite. Further, we generate a computer model based on our results which successfully recapitulate our experimental observations and creates new predictions that could broaden our understanding of root growth mechanisms in the dynamic environment.

## Introduction

The ability to sense and adapt to fluctuations in nutrient availability is essential for the survival of all organisms. Every life form on our planet possesses delicate mechanisms for sensing and reacting to the variable nutrient status and adjusts their behavior to maintain growth or cope with stress caused by malnutrition. Mineral nutrients absorbed from the soil are major determinants of plant growth and development. Although required, fluctuations in their availabilities either to sub- or supra-optimal levels often have detrimental effects on plant metabolism and physiology, thereby attenuating plant fitness. Hence, the acquisition of mineral nutrients from the soil needs to be tightly controlled and endogenous levels within a plant body maintained at a physiological optimum level. At the molecular level, balancing nutrient acquisition with the plant’s requirements implies that there is close communication between pathways controlling uptake, distribution and homeostasis of nutrients and the pathways coordinating plant growth and development.

The root system perceives and integrates local and systemic signals on the nutrient status to regulate activity of pathways mediating nutrient uptake and distribution. An important component of the plant’s nutrient management strategy involves a rapid modulation of the root growth and development. In response to nutrient availability, root meristem activity and elongation growth of primary root, as well as root branching, are adjusted in order to optimize nutrient provision to the plant body^1^. Production of new cells is essential for sustainable root growth; however, enhancement of the cell division machinery typically occurs within a range of hours^2^. In contrast, rapid modulation of cell elongation and manifold increase in cell volume would ensure faster growth responses^3^. Hence, in fluctuating environmental conditions root growth kinetics relies on the coordination of rapid elongation growth and adjustment of proliferation activity of the meristem.

Nitrogen (N) is a key macronutrient present in many key biological molecules and therefore constitutes a limiting factor in agricultural systems^4^. Although plants are dependent on an exogenous N supply and use nitrate (NO_3_^-^), nitrite (NO_2_^-^), and ammonium (NH_4_^+^) as major sources of inorganic N, their preference for different inorganic forms depends on plant adaptation to soil^4,5^. For example; wheat, maize, canola, beans, sugar beet, *Arabidopsis* and tobacco grow preferentially on NO_3_^−^ nutrition, whereas, rice and pine grow on NH_4_^+^ nutrition. Fluctuations in both concentrations and the form of nitrogen sources available in the soil have prominent effects on root system growth and development^6,7^. Deficiency in nitrogen severely interferes with root elongation growth and development; low to medium availability of nitrogen enhances root growth and branching to promote the exploitation of this macronutrient, whereas high levels of availability might inhibit the elongation growth of primary and lateral roots^8^. When exposed to local nitrate-rich zones, the root system responds by enhancing lateral root (LR) outgrowth^9–11^. In the model plant *Arabidopsis thaliana*, the local availability of NO_3_^−^ and NH_4_^+^ seems to have complementary effects on the LR development (NH_4_^+^ stimulates branching, whereas NO_3_^−^ induces LR elongation^11,12^). These complex adaptive responses of the root organ to N sources and heterogeneity in availability are regulated by a combination of systemic and local signaling^13^. The impact of available sources of N on the root system is closely interconnected with the activity of plant hormones including auxin, cytokinin, ABA, ethylene and others^14–16^. In recent years, a number of studies have demonstrated that auxin biosynthesis, transport, and accumulation is altered in response to different N regimes in maize^17,18^, soybean^19^, pineapple^20^ and *Arabidopsis thaliana*^16,21–23^. In *Arabidopsis*, several key auxin-related regulatory modules that respond to nitrogen availability were identified including *TAR2*, a gene involved in auxin biosynthesis, transporters of auxin such as *PIN-FORMED 1 (PIN1), PIN2, PIN4* and *PIN7* and molecular components, which control their subcellular trafficking^21,24^. At the level of auxin signaling, Auxin Response Factor *AUXIN RESPONSE FACTOR 8* (*ARF8*, encoding a transcription factor of the auxin signaling machinery) was identified as a N responsive gene in the pericycle^25^. *ARF8* together with its associated microRNA167s is involved in the control of the ratio between LR initiation and emergence^25–28^. Another mechanism of nitrogen – auxin interplay underlying adaptation of the root system is mediated through NRT1.1, nitrate transceptor ^29^. Its dual auxin-nitrate transport activity has been shown to play an important role in the adaptation of the root system, in particular, LR emergence to nitrate availability^21,30^.

Flexible modulation of primary root growth to fluctuations in nitrogen resources has been recognized as a prominent foraging strategy to optimize N exploitation^31^. However, the mechanisms that control the rapid reconfiguration of root growth dynamics in response to diverse N sources are still poorly understood. Here, to dissect the tissue and cellular mechanisms underlying the early phases of this adaptive process we focused on the primary responses of *Arabidopsis* roots to alterations in the available source of N such as NH_4_^+^ and NO_3_^-^. We performed real time vertical confocal imaging to capture the earliest root responses after the replacement of NH_4_^+^ by NO_3_^-^. We found that in roots supplied with NH_4_^+^, local attenuation of meristematic activity in the epidermis results in the earlier transition of epidermal cells into elongation when compared to the cortex, thus generating asynchronous elongation of the adjacent tissues. Substitution of NH_4_^+^ for NO_3_^-^ led to a rapid enhancement of root growth associated with the simultaneous entrance of more cells at the root transition zone into elongation, and the subsequent re-establishment of a critical balance between cell proliferation and elongation. We demonstrate that root epidermis and cortex tissues supplemented with NO_3_^-^ synchronize their growth patterns. We show that the essential mechanism underlying this flexible adaptation of root growth involves nitrate-dependent fine-tuning of the auxin transport mediated by PIN2. In roots supplied with different forms of N, distinct localization patterns of PIN2 are generated as a result of dynamic PIN2 subcellular trafficking. Intriguingly, phosphoproteome analysis of PIN2 (Vega et al.) led to the identification of an uncharacterized nitrate-sensitive phosphorylation site. The functional characterization of PIN2 and its phosphor-variants suggest that the N source dependent modulation of PIN2 phosphorylation status has a direct impact on the flexible adjustment of PIN2 localization pattern, and thereby facilitates the adaptation of root growth to varying forms of N supply. Finally, we integrated experimental data regarding the nitrogen-dependent root growth into a quantitative computer model. Our computer model recapitulated *in planta* patterning from a minimal set of assumptions and made predictions that were tested experimentally. Taken together, we present a quantitative mechanistic model of how *Arabidopsis* primary root growth is fine-tuned to different N sources. We hypothesize, that the flexible modulation of growth patterns relying on nutrient response on auxin transport is an important part of the intelligent strategy, to enable plant root adaption to the dynamically changing environment and thus maintain its sustainable growth.

## Results

### Root growth rapidly adjusts to form of nitrogen source

To explore how primary root responds and adapts to different forms of N, *Arabidopsis* seedlings were grown on NH_4_^+^ as an exclusive N source for five days (5 DAG) and afterward transferred on media containing either NH_4_^+^ or NO_3_^-^. We found that replacement of NH_4_^+^ by NO_3_^-^ rapidly enhanced root length and already 6 hours after transfer (HAT), roots were significantly longer compared to these supplied with NH_4_^+^ (Fig. S1a). In general, root growth is determined by the elongation of cells, which are constantly produced by the root apical meristem. To study processes that underlie the adaptation of root growth to different forms of N a vertical confocal microscope equipped with a root tracker system was employed. Using this setup, we were able to detect and monitor the earliest root responses with a high cellular resolution^32^. To minimize the interference of physiological conditions for seedling development, a light-dark regime was maintained in course of the root tracking. After the transfer of wild type (Col-0) seedlings to NH_4_^+^ containing medium root growth rate (RGR) was enhanced, presumably as a response to stress caused by transfer of seedlings to a fresh plate. Within ∼120 min RGR stabilized at an average speed of 1.37 ± 0.025 µmmin^-1^. Transition to dark period correlated with a rapid drop of RGR to 0.98 ± 0.029 µmmin^-1^, which was maintained during the dark phase and at the light recovered again to 1.27 ± 0.048 µmmin^-1^. Seedlings transferred to NO_3_^-^ reacted by an increase of RGR to 1.77 ± 0.042 µmmin^-1^ and similarly to roots on NH_4_^+^, during the dark period their RGR decelerated and was retrieved to 1.81 ± 0.051 µmmin^-1^ at the light (Fig. 1a, Supplemental video 1). Hence, provision of NO_3_^-^ caused a rapid enhancement of RGR when compared to NH_4_^+^, but it did not interfere with its circadian rhythmicity^33^.

**Figure 1.**
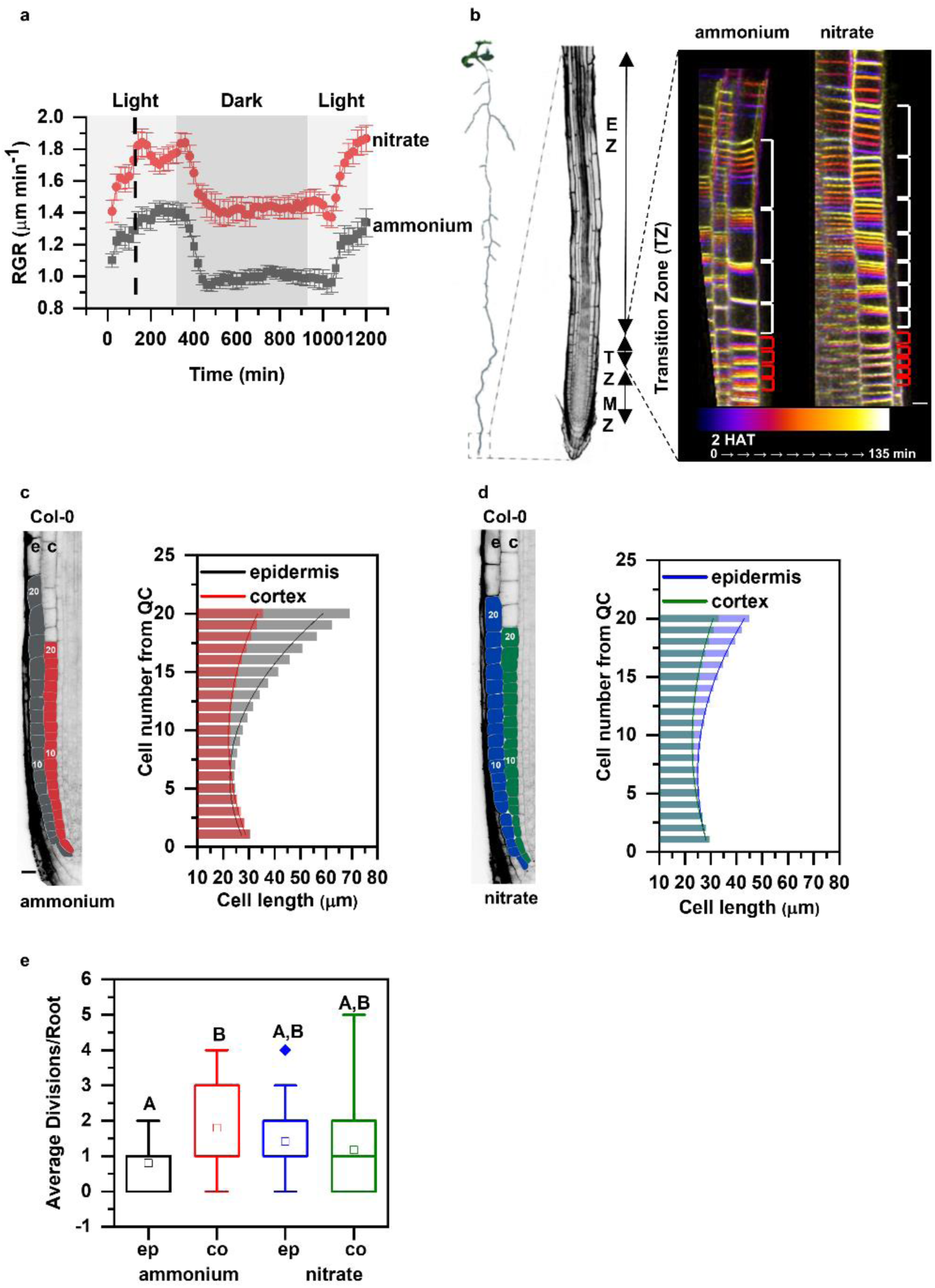
Primary root growth kinetics of Arabidopsis (*Arabidopsis thaliana (L.) Heynh. Columbia-0*, Col-0) on ammonium or nitrate containing medium. **a**. Seedlings were transferred 5 days after germination (DAG) to medium supplemented with ammonium (grey) or nitrate (red). Root growth rates (RGR in µm/min) were monitored over a 1200 minutes period. Data represent the geometric mean (± standard error, SE) of three independent experiments (each consisting of 3 roots per treatment). Light and dark periods are indicated as light or dark gray background, respectively. **b**. On the left, schematic representation of distinct root zones: Meristematic Zone (MZ), Transition Zone (TZ) and Elongation Zone (EZ). On the right, time lapse imaging of cell growth at the TZ. Cells were visualized using the plasma membrane marker (wave line W131Y). Observation of roots started 2 hours after transfer (2 HAT; blue) on ammonium or on nitrate for 135 min (white) and images were recorded every 20 minutes (9 stacks/root/recording). Red and white brackets indicate the length of meristematic and elongating cells at the last measurement point, respectively. Scale bar = 30 µm. **c and d**. Representation and quantification of cell length in epidermal (e) and cortical (c) cell files. Optical, longitudinal sections of 5 DAG old Col-0 roots 12 HAT to ammonium (C) or nitrate (D) supplemented media. The first 20 epidermal (e) and cortex (c) cells (from quiescent center (QC)) are highlighted in grey and in red on ammonium (C), and in blue and green on nitrate (D), respectively. Scale bar = 30 µm. Column bars denote the geometric mean of the cell lengths at the respective positions. Lines represent a polynomial regression fit, with calculated slopes between cells 10 and 20 of 3.32639±0.17172 (ammonium, epidermis), 1.22033±0.08754 (ammonium, cortex) and 1.70502±0.09532 (nitrate, epidermis), 0.82342±0.06973 (nitrate, cortex). Data are derived from 3 independent experiments; total number of analyzed roots are n=18 in each case. **e**. Graphical representation of the average number of cell divisions along epidermis (ep) and cortex (co) in 5 DAG root tips 12 HAT to ammonium or nitrate supplemented media. Data are derived from 15 and 17 roots. The statistical significance was evaluated with ANOVA at p<0.05.

To gain more insight into the mechanistic basis underlying the rapid increase of root length after substitution of NH_4_^+^ for NO_3_^-^, we focused on cells in the transition zone (TZ). The TZ is located between the root apical meristem and elongation zone, and cells while passing this developmental zone undergo essential modifications associated with their transition from the proliferative to the elongation phase^34,35^ (Fig. 1b). Time-lapse experiments capturing root growth from 2 to 3.76 hours after transfer combined with a tracking of cell membranes pointed at differences in the elongation pattern of epidermal cells in roots supplied with either NH_4_^+^ or NO_3_^-^. While, in roots supplemented with NH_4_^+^ only a few epidermal cells enter into elongation phase. Provision of NO_3_^-^ increased number of elongating cells in the TZ (Fig. 1b, Supplemental video 2). Next, we analyzed in detail 18 roots 12 HAT on either NH_4_^+^ or NO_3_^-^ and measured length of the epidermal cells across the meristematic, transition and the start of the elongation zones. The analyses suggested that on NO_3_^-^ more epidermal cells enter into transition phase, as indicated by an increased number of cells 30-40 μm long when compared to roots on NH_4_^+^ (Fig. S1b). Despite the stimulating impact of NO_3_^-^ on cell transition into the elongation phase, no differences in the maximal length of fully differentiated epidermal cells between roots on NO_3_^-^ and NH_4_^+^ were detected (Fig. S1c). This suggests that NO_3_^-^ promoted root growth is a result of modulated elongation kinetics of cells along the longitudinal root growth axis and not increase of the maximal cell length.

To sustain root growth, the rate of cell elongation and differentiation has to be tightly balanced with the production of new cells in the root meristem^36^. Hence, enhanced growth of cells after replacement of NH_4_^+^ by NO_3_^-^ could lead to depletion of the meristem if expansion of cells would prevail over a new cell production. To examine how root meristem adapt to change in N supply, cell length and frequency of divisions in epidermis and cortex along the longitudinal root growth axis were closely inspected 12 HAT. Surprisingly, length of epidermal cells started to increase from the 11^th^ cell on (cell number was counted from quiescent center (QC)) in roots supplied with NH_4_^+^ (Fig. 1c and Supplemental Document 1a-b). In contrast, roots on NO_3_^-^ exhibited an increase in size from the 13^th^ epidermal cells (Fig. 1d, S1d Supplemental Document 1a-b). Unlike the epidermis, the length profiles of cortex cells were not significantly different between roots supplied with either NH_4_^+^ or NO_3_^-^ (Fig. S1e, Supplemental Document 1a-b). Therefore, the growth of epidermal and cortex cells in roots on NH_4_^+^ displayed clearly asynchronous behavior (Fig. 1c, d). Additionally, a machine learning approach was applied to regression analysis for assessing the importance of each variable (i.e. treatments: ammonium and nitrate, tissues: epidermis and cortex, cell positions) on cell length differences. Analysis of deviance was followed by estimated marginal mean (emmean) comparisons of cell lengths in different tissues (epidermis vs cortex) at each cell position (1-20 from QC) for each treatment (ammonium vs nitrate). The results show that ammonium and nitrate treatments affect the cell positions differentially: epidermal cells from the 17th up to the 20th position are significantly longer on ammonium while cell length in cortex is not affected by the treatments. (Supplemental Document 1a-b). Results were confirmed by recursive partitioning analysis and shown in a decision tree (Supplemental Document 1c).

The distinct elongation pattern of epidermal and cortex cells detected in roots on NH_4_^+^ can only be sustained if cell divisions in cortex compensate for an earlier start of cell elongation in the epidermis. Accordingly, the scoring of cell division events (visualized by DAPI) revealed a higher number of mitotic events in cortex compared to epidermal cells in roots transferred to NH_4_^+^. On NO_3_^-^ similar frequency of cell divisions in both epidermal and cortex cell files was observed (Fig. 1e, Fig. S2a). Finally, monitoring of the cell cycle reporter *CyclinB::GUS* expression 2 days after transfer (DAT) to either NH_4_^+^ or NO_3_^-^ revealed enhanced reporter expression and overall enlargement of the meristematic zone in roots supplemented with NO_3_^-^ (Fig. S2b).

Altogether, these data indicate that roots adopt distinct growth strategies involving fine-tuning of cell division and expansion across adjacent tissues to adapt to different forms of N. In roots supplied with NH_4_^+^, the meristematic activity of epidermal cells is attenuated, which results in their earlier transition into the elongation phase when compared to the cortex. Provision of NO_3_^-^ increases the number of epidermal cells in the TZ (Fig. 1b, S1b), which is one of the earliest detectable adaptive responses. Subsequently, within twelve hours, the frequency of cell division in the epidermis increases, which results in shift of balance between cell division and elongation and more synchronized growth of cortex and epidermis. Eventually, a long-term supply of NO_3_^-^ enables enlargement of the root apical meristem compared to roots supplied with NH_4_^+^.

### Level and pattern of auxin activity in roots are modulated by form of nitrogen source

The plant hormone auxin is an essential endogenous regulatory cue that determines key aspects of root growth. Interference with auxin biosynthesis^37^, signaling^38^ or distribution^39^ at the root tip has a significant impact on the meristem maintenance, and transition of meristematic cells into elongation and differentiation phase. Distinct growth patterns observed in roots supplemented with different forms of N prompted us to monitor distribution of auxin at the root tip. Quantification of the LUCIFERASE activity in protein extracts from roots carrying the auxin sensitive *DR5::LUCIFERASE* reporter revealed that already one hour after transfer to NO_3_^-^ containing medium auxin response increases when compared to roots transferred to NH_4_^+^ supplemented medium (Fig. S3a). To closely inspect the auxin distribution in a cell lineage-specific manner a ratiometric degradation based *R2D2* auxin reporter was implemented^40^. In accordance with observations based on the *DR5::LUCIFERASE* reporter, a decreased ratio between DII-Venus (green) and mDII-Tomato (red) fluorescent signals indicated increased levels of auxin activity in the central cylinder of roots in response to replacement of NH_4_^+^ by NO_3_^-^ (Fig. S3b).

In addition, we focused on the detailed profiling of the R2D2 reporter in the epidermis and the cortex (Fig. S3c). Interestingly, we detected an overall increase of auxin activity in epidermal cells when compared to cortex cells in roots supplied with NH_4_^+^, whereas no difference between these two cell files were detected in roots on NO_3_^-^ (Fig. 2a, b). Furthermore, on NH_4_^+^ there was an increase of auxin activity in epidermal cells when compared to cortex cells (starting from ∼ 11^th^ cell from the QC), while in roots supplied with NO_3_^-^ the auxin activity profiles followed similar trends of steady increase in both cortex and epidermal cell files (Fig. 2a, b). Altogether, these analyses indicate that pattern of auxin activity at root meristems might adapt to specific N conditions. In roots supplied with NH_4_^+^, the early steep gradient of auxin signaling in epidermal cells correlates with their early transition into the rapid elongation phase. Whereas in cortex cells, auxin reaches concentrations which might drive elongation in more proximal cells. Substitution of NH_4_^+^ by NO_3_^-^ attenuates differences in profiles of the auxin distribution between the cortex and the epidermal cell files, which would lead to the synchronized cell growth (Fig. 2a, b compared to Fig. 1c, d).

**Figure 2.**
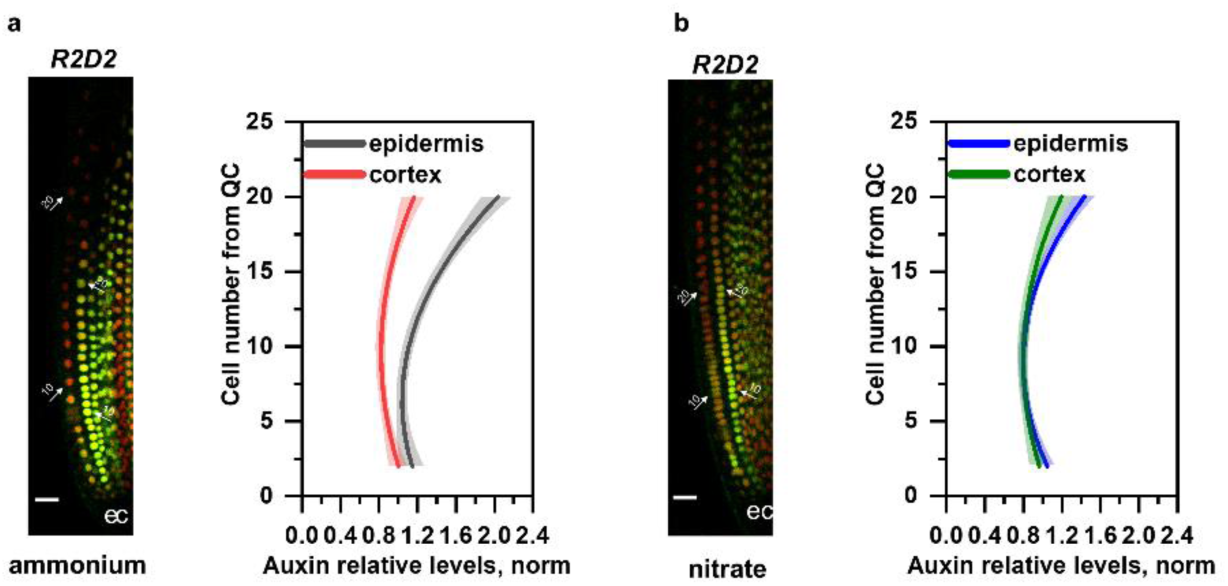
Relative auxin level in Col-0 root tips transferred to medium supplemented with ammonium or nitrate. Maximum intensity Z-stack projection images of 5 DAG old roots expressing the *R2D2* auxin signaling reporter 12 HAT to ammonium (**a**) or nitrate (**b**) supplemented media. White arrows mark the position of the 10^th^ and 20^th^ cells from QC; “e” and “c” mark epidermis and cortex, respectively. Scale bar = 50 µm. Graphs denote normalized relative auxin levels at the respective positions. Lines represent polynomial regression fit with 95% confidence band. Data are derived from 5 roots per condition from three independent experiments.

### Nitrogen source affects basipetal auxin transport

Directional cell-to-cell transport of auxin significantly contributes to the estalishment of the auxin activity pattern at the root tip. The Polar auxin transport (PAT) machinery, composed of AUX/LAX influx and PIN efflux carriers, directs the flow of auxin from the shoot acropetally through the stele towards the root tip; from where it is via epidermis basipetally redistributed to the elongation zone. At the TZ, auxin might be redirected from the basipetal stream across the cortex, endodermis and pericycle back to stele and root tip, thereby fine-tuning levels of auxin at the TZ^41,42^. The modulation of auxin activity pattern in the outer tissues detected after the replacement of NH_4_^+^ for NO_3_^-^ suggests that there are alterations of the basipetal auxin transport. To explore how different forms of N affect the flow of auxin in basipetal direction, transport assays using radioactively labeled auxin (^3^H-IAA) were performed. Six hours after applying ^3^H-IAA to the root tip, radioactivity in the proximal zone of the primary roots supplied with NH_4_^+^ was significantly lower when compared to roots on either NO_3_^-^ supplemented or standard Murashige and Skoog (MS) medium (Fig. 3A). These results indicate that basipetal auxin transport can be modulated by available source of N, and provision of NO_3_^-^ enhances flux of auxin in shootward direction when compared to NH_4_^+^.

**Figure 3.**
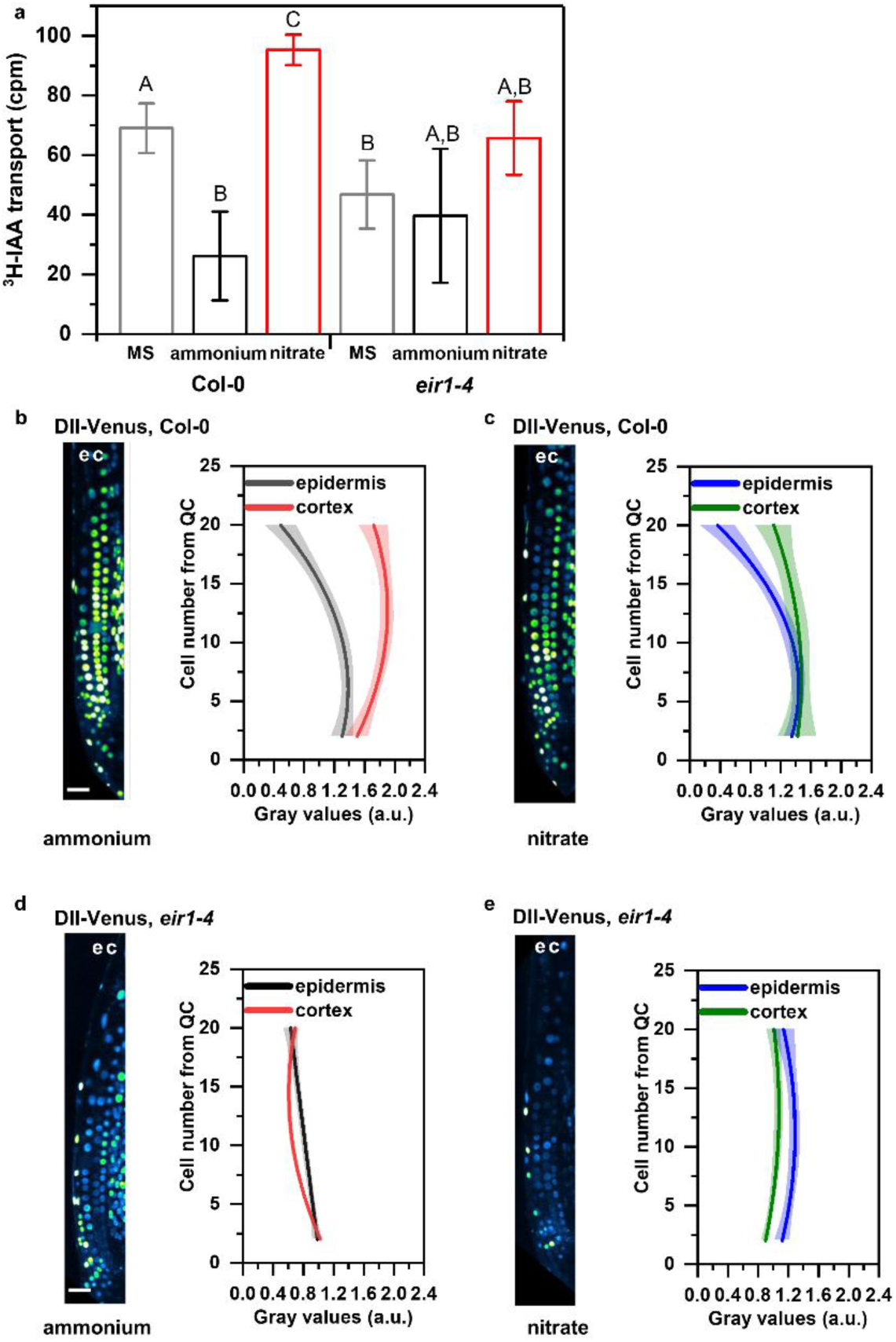
Monitoring basipetal auxin transport and auxin response at root tips of Col-0 and *eir1-4* roots transferred to ammonium or nitrate containing medium. **a**. Basipetal (shootward) auxin transport measurements in Col-0 and *eir1-4* roots grown on control Murashige and Skoog (MS) or with either nitrate or ammonium supplied media. ^3^H-IAA was applied at the root tip of 7 DAG wild-type (Col-0) or *eir1-4* seedlings. Radioactivity was measured 6 h after application of ^3^H-IAA in root segments after excision of the apical ≈ 1 mm of the root tip. Values shown are the geometric mean (± standard error, SE) for at least 30 seedlings. The amount of auxin transported into each root segment for Col-0 and *eir1-4* was compared by ANOVA at p<0.05. cpm, counts per minute. **b-e**. Maximum intensity Z-stack projection images of 5 DAG old Col-0 and *eir1-4* roots expressing the *DII-Venus* auxin signaling reporter 12 HAT to ammonium (**b and d**) or nitrate (**c and e**) supplemented media. “e” and “c” mark epidermis and cortex, respectively. Scale bar = 50 µm. Graphs denote normalized relative auxin levels at the respective positions. Lines represent polynomial regression fit with 95% confidence band. Data are derived from measurements of n=8 (ammonium) and n=10 (nitrate) roots of Col-0 and n=10 roots of *eir1-4* per condition.

The PIN2 auxin efflux carrier is amongst the principal components of PAT mediating basipetal transport of auxin in roots^43,44^. To test whether adjustment of the basipetal auxin flow in response to different sources of nitrogen is dependent on activity of PIN2, we tested *eir1-4*, a mutant defective in this efflux transporter. In agreement with previous reports^45^, a significantly lower radioactivity in the proximal root zone of the *eir1-4* was detected when compared to wild type roots on MS medium (Fig. 3a). Noteworthy, no radioactivity increase in the proximal zone of *eir1-4* roots was observed in roots supplied with NO_3_^-^ when compared to NH_4_^+^ (Fig. 3a), pointing towards PIN2 function in the flexible adjustment of the basipetal auxin flow in response to form of N source. To further examine the role of the PIN2 mediated transport in establishment of distinct auxin patterns at root tips supplemented with different forms of N, we monitored the auxin sensitive reporter DII-Venus and its stabilized auxin responsive analog mDII-Venus^46^ as a reference in *eir1-4* and Col-0 roots. The expression pattern of DII-Venus reporter in Col-0 roots was largely consistent with what we observed using the R2D2 reporter (Fig. S4a-b). In Col-0 roots supplied with NH_4_^+^, a reduced DII-Venus signal indicated a higher auxin activity in epidermal cells when compared to the cortex. Also, consistently with the R2D2 reporter, a steeper slope of auxin activity in epidermis when compared to cortex (with onset at ∼8^th^ cell distance from QC) was detected in roots supplied with NH_4_^+^, whereas in roots on NO_3_^-^, auxin activity both in epidermis and cortex followed similar trends (Fig. 3b, c compared to Fig. 2a, b and Fig. S4a, b). *eir1-4* was severely affected in adjustment of auxin pattern to different N sources. When compared to Col-0, overall higher levels of auxin activity in both epidermal and cortex cells and a shallower slope of auxin activity increase in the epidermis was observed in *eir1-4* roots supplied with NH_4_^+^. As a result, the difference in auxin activity profiles between the cortex and the epidermis in *eir1-4* was less pronounced than in wild type roots (Fig. 3d compared to Fig. 3b and Fig. S4a). On NO_3_^-^, overall profiles of auxin activity in epidermis and cortex of *eir1-4* followed similar trends, characterized by shallow slope along the longitudinal root growth axis (Fig. 3e, Fig. S4b). Importantly, expression pattern of the auxin insensitive mDII-Venus reference construct remained largely unchanged under all tested conditions in both wild type and *eir1-4* (Fig. S4c, d). Altogether, our results point at an important role of PIN2 dependent basipetal auxin transport in adjustment of auxin activity pattern in roots to specific N conditions.

### PIN2 mediates root growth adaptation to nitrogen resources

To further examine the role of PIN2 mediated basipetal auxin transport in root growth adaptation to different sources of N, *eir1-4* and *eir1-1* mutant alleles of *PIN2* were analyzed. Unlike in wild type, no significant increase in root length was detected 1 DAT in either *eir1-4* or *eir1-1* seedlings on NO_3_^-^ when compared to NH_4_^+^ supplemented medium (Fig. S5a). Closer inspection of the RGR in real time using vertical confocal - root tracking set up showed that after transfer on NH_4_^+^ growth of the *eir1-4* roots stabilized at 1.47 ± 0.041 µmmin^-1^ and 1.35 ± µmmin^-1^ during light and dark period, respectively. However, no significant increase of RGR after transfer to NO_3_^-^ containing medium could be observed (Fig. 4a). These results strongly support an essential role of PIN2 mediated basipetal auxin transport in rapid adjustment of root growth to form of nitrogen source.

**Figure 4.**
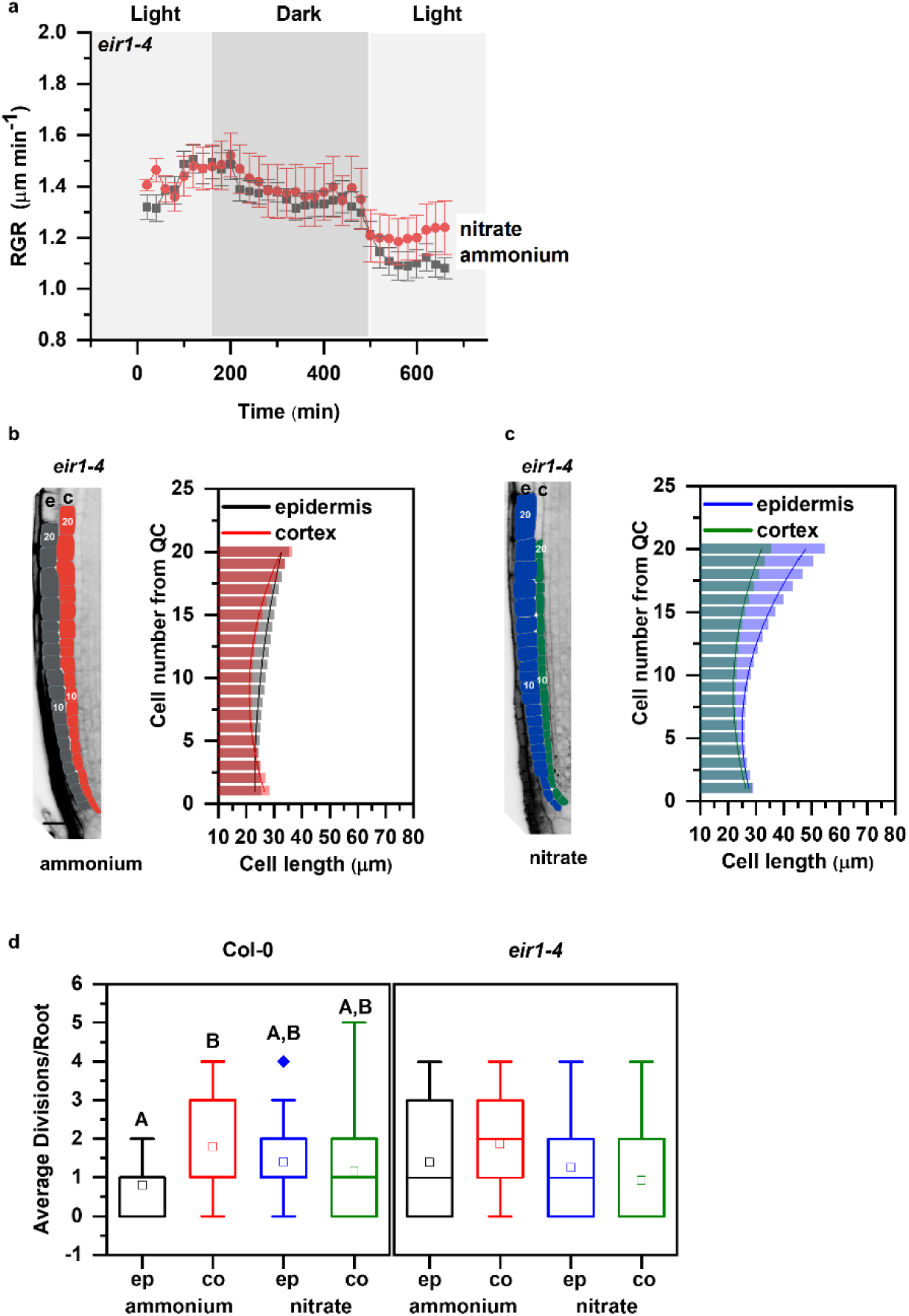
Primary root growth kinetics of *eir1-4* roots transferred to ammonium or nitrate amended medium. **a**. Root growth rate (RGR in µm/min) of *eir1-4* roots transferred 5 DAG to ammonium (grey) or nitrate (red) containing medium over a period of 680 minutes. Data represent the geometric mean (± standard error, SE) of three independent experiments (number of roots n=5 in each case). Light and dark periods are highlighted in light or dark gray. **b and c**. Representation and quantification of cell length in epidermal (e) and cortical (c) cell files. Optical, longitudinal sections of 5 DAG *eir1-4* roots 12 HAT to ammonium (B) or nitrate (C) supplemented media. The first 20-20 epidermal and cortex cells (from quiescent center (QC)) are highlighted in grey and in red on ammonium (B) and in blue and green on nitrate (C), respectively. Scale bar = 30 µm. Column bars denote the geometric mean of cell length at the respective positions. Lines represent a polynomial regression fit, with calculated slopes between cells 10 and 20 of 0.75884±0.02624 (ammonium, epidermis), 1.13088±0.08446 (ammonium, cortex) and 2.06912±0.10341 (nitrate, epidermis), 0.99878±0.07278 (nitrate, cortex). Data are derived from 3 independent experiments, total number of the analyzed roots are n=9, ammonium and n=8, nitrate. **d**. Average number of cell divisions along the epidermis (ep) and cortex (co) in 5 DAG old Col-0 and *eir1-4* root tips 12 HAT to ammonium or nitrate supplemented media. Data are derived from n=15 and n=17 roots of Col and n=10 and n=9 roots of *eir1-4* on ammonium and nitrate, respectively. Statistical significance was evaluated with ANOVA at p<0.05.

To explore whether *eir1-4* root growth adapts to different forms of N, elongation patterns of epidermal and cortex cells were analyzed. Measurements of cell lengths along the longitudinal growth axis of *eir1-4* roots supplied with NH_4_^+^ revealed that unlike in Col-0, epidermal cells undergo gradual, steady elongation growth comparable to that in cortex. Notably, patterns of cortex and epidermal cell growth in *eir1-4* appear more synchronous than in wild-type roots on NH_4_^+^ (Fig. 4b versus Fig.1c). In *eir1-4* roots 12 HAT from NH_4_^+^ to NO_3_^-^ supplemented medium we observed largely synchronized pattern of elongation in both epidermal and cortex cell files, characterized by gradual, steady increase of cell length similar to these observed in Col-0 (Fig. 4c and Fig. 1d). Consistently with a more synchronous pattern of epidermal and cortex cell growth in both N regimes, no significant differences in frequency of mitotic events between epidermis and cortex were found in *eir1-4* roots on medium supplied with either NH_4_^+^ or NO_3_^-^ (Fig. 4d).

Overall, loss of PIN2 activity interfered with enhancement of root growth in response to NO_3_^-^ provision and affected the establishment of tissue specific growth patterns typically adopted by Col-0 roots supplied with different sources of N. Altogether, these results indicate that PIN2 mediated basipetal auxin transport plays an important function in acquiring distinct root growth patterns during adaptation to different N sources.

### PIN2 delivery to the plasma membrane and polarity is adjusted in response to form of nitrogen source

To explore the mechanisms underlying PIN2 function in root growth adaptation to different N sources we examined its expression, abundance at the plasma membrane (PM) and subcellular trafficking in roots supplied with NH_4_^+^ or NO_3_^-^. RT-qPCR analyses of 7 DAG roots grown on NH_4_^+^ and transferred to media supplemented with either NH_4_^+^ or NO_3_^-^ for 1, 6 and 48 hours did not reveal any significant changes in *PIN2* transcription in any of the tested conditions (Fig. S6a). Likewise, expression of neither the *PIN2::nlsGFP* nor the *PIN2::GUS* reporter was affected by different N source (Fig. S6b). Interestingly, monitoring of *PIN2::PIN2-GFP* transgenic seedlings revealed significantly increased abundance of the PM located PIN2-GFP in epidermal and cortex cells of roots supplied with NO_3_^-^ when compared to NH_4_^+^ (Fig. 5a). Furthermore, in cortex cells at the transition zone of NO_3_^-^ supplied roots, besides expected localization at the apical PM^39^, enhanced lateralization of PIN2-GFP to the inner and outer PMs could be detected (Fig. 5b, Fig. S6d). Immunolocalisation using PIN2-specific antibodies is fully consistent with the observations of PIN2-GFP and ruled out possible interference with fluorescence of GFP reporter by different N source (Fig. S7a-c). Hence, substitution of NH_4_^+^ by NO_3_^-^ seems to affect PIN2 at post-transcriptional rather than at transcriptional level.

**Figure 5.**
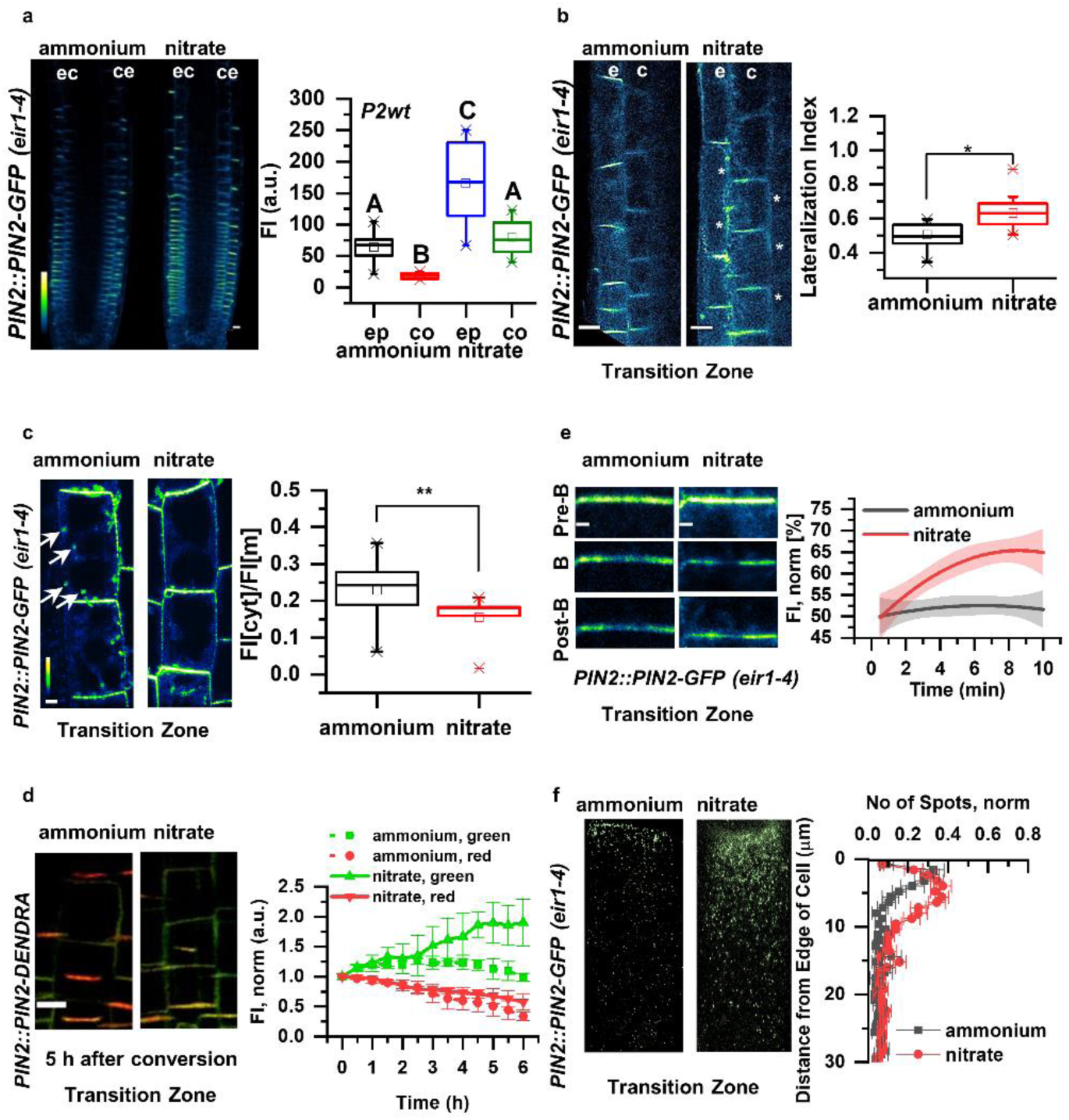
PIN2 protein abundance, polarity and vacuolar trafficking in roots on ammonium or nitrate containing medium. **a**. Pseudo-colored, optical longitudinal cross sections of 5 DAG roots expressing *PIN2::PIN2-GFP, eir1-4* 12 HAT to ammonium or nitrate supplemented media. “e” denotes epidermis and “c” cortex, respectively. Color code represents GFP intensity from low (blue) to high (white) values. Scale bar = 20 µm. Box plots display the distribution of the cell membrane derived PIN2-GFP fluorescence intensity (FI) values (in arbitrary units, a.u.) on ammonium (grey, epidermis (ep) and red, cortex (co), n=15) and nitrate (blue, epidermis (ep) and green, cortex (co), n=10) grown roots. 5 cells per root were analyzed. The statistical significance was evaluated with ANOVA at p<0.05. **b**. Higher magnification of pseudo-colored confocal images of 5 DAG old roots expressing PIN2-GFP 12 HAT to ammonium or nitrate supplemented media. “e” denotes epidermis and “c” cortex, respectively. Color code represents GFP intensity from low (blue) to high (white) values. Scale bar = 12 µm. White stars mark PIN2-GFP protein localization on the lateral membranes. Box plots display lateralization index (fluorescent signal detected on apical/basal membranes divided by the signal value at inner/outer membranes) of roots on ammonium (n=31 cells from 6 roots) or nitrate (n=24 cells from 6 roots) supplemented medium. **c**. Pseudo-colored PIN2-GFP signal in epidermal cells of 5 DAG old roots 12 HAT to ammonium or nitrate containing media. White arrows point to PIN2-GFP containing intracellular vesicles. Box plots represent the ratio in fluorescent signal detected inside the cell vs on the membranes (FI[cyt]/FI[m]). n=6 roots per condition, 5 cells per root analyzed. Scale bar = 5µm. **e**. FRAP analysis of PIN2 protein mobility in *PIN2::PIN2-GFP* expressing epidermal cells 12 HAT to ammonium or nitrate. The graph shows polynomial regression fit with 95% confidence band of the mean signal recovery in the bleached region of interest (ROI) after background subtraction and normalization to photobleaching. Data are derived from 3 independent experiments, each consisting of 5 membranes from 3 different roots. Scale bar = 2 µm. **d**. Microscopic images showing PIN2-Dendra fluorescent signal five hours after photoconversion of PIN2-Dendra into its red form. Depletion of the red signal and recovery of the green signal over a 6 hours period was followed in parallel in 5 DAG old roots12 HAT to ammonium or nitrate supplemented media. Note the increase in the intensity of the green signal in roots transferred to nitrate. Graph represents the mean signal ± SD (n=6 roots per condition, 20 cells per root analyzed). The experiment was repeated 3 times. Scale bar = 20 µm. **f**. Representative 3D SIM microscopic images of 10 DAG old epidermal cells expressing *PIN2-GFP* 12 HAT to ammonium or nitrate containing media. Green dots represent PIN2-GFP on the lateral cell surface (polar domain) of epidermal cells in the transition zone. Graph represents the number of GFP positive spots along a 30 µm long region starting at the apical side of the cell (8 cells per 4 roots and 9 cells per 4 roots) were analyzed per treatment, experiment was done 3 times. Note the effect of ammonium versus nitrate on the distribution of the PIN2-GFP spots.

PIN proteins constantly recycle between the PM and endosomal compartments, thus their abundance at the PM is largely dependent on a balance between endo- and exocytosis^47,48^. Hence, we explored whether modulation of PIN2 subcellular trafficking is the mechanism involved in adjustment of the PIN2 pattern in response to the available N source. In epidermal cells on NH_4_^+^ when compared to NO_3_^-^ supplied roots, the ratio between intracellular versus PM-located PIN2-GFP was shifted in favor of intracellular localization and frequently endosomal vesicles with PIN2-GFP signal could be detected (Fig. 5c, Supplemental video 3). This indicates that dynamics of PIN2 subcellular trafficking might be altered on the basis of the N source. To assess whether in NH_4_^+^ versus NO_3_^-^ supplied roots, accumulation of PIN2 at the PM is the result of a changed balance between endo- and exocytosis, we analyzed *pPIN2::PIN2-Dendra* seedlings. The irreversible photo-conversion of the Dendra fluorochrome by UV light from its green form to red allowed us to follow the impact of the N source on the subcellular fate of PIN2. By monitoring the PIN2-Dendra signal after photo-conversion (red signal) versus the newly synthesized PIN2-Dendra (green signal) in real time we could evaluate the kinetics of PIN2 internalization from the PM and delivery of the *de novo* synthesized PIN2-Dendra proteins. We found that the kinetics of the photo-converted PIN2-Dendra (red signal) at the PM in either NH_4_^+^ or NO_3_^-^ were not statistically different, indicating that the internalization of PIN2 is not affected by the N source. Nevertheless, recovery of the newly synthesized PIN2-Dendra (green signal) was significantly enhanced in NO_3_^-^ when compared to NH_4_^+^ supplied roots (Fig. 5d, Fig. S6c). Considering that different sources of N did not have significant impact on *PIN2* transcription (Fig. S6a), these results suggest that recycling or secretion of PIN2 to the PM is more promoted in NO_3_^-^ supplied roots than in those on NH_4_^+^. To further examine the impact of N source on the delivery of PIN2 to the PM we performed Fluorescence Recovery After Photobleaching (FRAP) analyses on the apical membrane of the cell. Lateral diffusion of PIN proteins at the PM is negligible^49^, thus PIN2-GFP signal recovery after photobleaching can be correlated with the delivery of PIN2 protein to the PM. In epidermal cells of NO_3_^-^ supplied roots PIN2-GFP signal recovered significantly faster as compared to roots supplied with NH_4_^+^ (Fig. 5e), thus strongly suggesting that delivery of PIN2 towards the PM is differentially regulated by specific forms of N source. Finally, to examine whether the above described different recycling behavior of PIN2 has an impact on the establishment of its apical polar domain, we performed super-resolution imaging employing three-dimensional structured illumination microscopy (3D-SIM). In roots supplemented with either NH_4_^+^ or NO_3_^-^, PIN2-GFP accumulated at the apical edge of epidermal cells to the same level. However, in NH_4_^+^ supplemented roots, number of the PIN2-GFP positive particles decreased with distance from the cell edge significantly more than in roots supplied with NO_3_^-^ (Fig. 5f).

In summary, these results suggest that PIN2 subcellular trafficking, and in particular the delivery of PIN2 to the PM is differentially adjusted according to the N source.

### Nitrogen dependent PIN2 phosphorylation fine-tunes intracellular dynamics and membrane polarity of PIN2

Post-translational modifications including phosphorylation are regulatory cues with significant impact on the intracellular trafficking and polar membrane localization of PIN proteins^50^. Phosphoproteome analysis of samples with either NH_4_^+^or NO_3_^-^ as their N source revealed that PIN2 was among the proteins exhibiting an altered pattern of phosphorylation in response to NH_4_^+^ (Vega et al.). Ser439 located at the very end of the PIN2 cytoplasmic loop (C-loop), was identified as a potential target for differential phosphorylation, where a reduction of phosphorylation in NO_3_^-^ conditions was detected compared root supplied with either NH_4_^+^ or KCl (Vega et al.). Multiple sequence alignment revealed that this Ser439 residue is highly specific to PIN2 (Fig. S6e). Interestingly, amino acid sequence alignment of PIN2 orthologues indicated that Ser439 is highly conserved in the PIN2 or PIN2-like clade across plant species including gymnosperms, mono- and dicotyledonous plants (Fig. S6f).

To examine a role of this specific, uncharacterized phosphosite in subcellular dynamics and function of PIN2, we introduced amino acid substitutions S439D and S439A to achieve either gain- or loss-of phosphorylation status of PIN2, respectively. *PIN2::PIN2*^*S439D*^*-GFP* and *PIN2::PIN2*^*S439A*^*-GFP* constructs were introgressed into the *eir1-4* mutant line. The phosphor-variant version PIN2^S439D^-GFP, like PIN2-GFP, accumulated at the PM of epidermal and cortex cells significantly more in roots supplied with NO_3_^-^ than NH4^+^ (Fig. 6a, b, d). Interestingly, the amount of the PM localized phospho-dead PIN2^S439A^-GFP in NH_4_^+^ -supplied roots was significantly higher when compared to PIN2-GFP and PIN2^S439D^-GFP, and only a slight increase in epidermal cells could be detected in response to NO_3_^-^ supply (Fig. 6a, c, d). Furthermore, in cortex cells at the TZ, reduced lateralization of PIN2^S439D^-GFP on NO_3_^-^ supplemented medium could be observed. PIN2^S439A^-GFP lateralized towards outer and inner PMs irrespective of the N source thus phenocopying the PIN2-GFP pattern in NO_3_^-^ supplied roots (Fig. 6e-g). Altogether, these results suggest that phosphorylation status of PIN2 on S439 account for fine-tuning of PIN2 trafficking towards the PM and polarity establishment under varying N sources.

**Figure 6.**
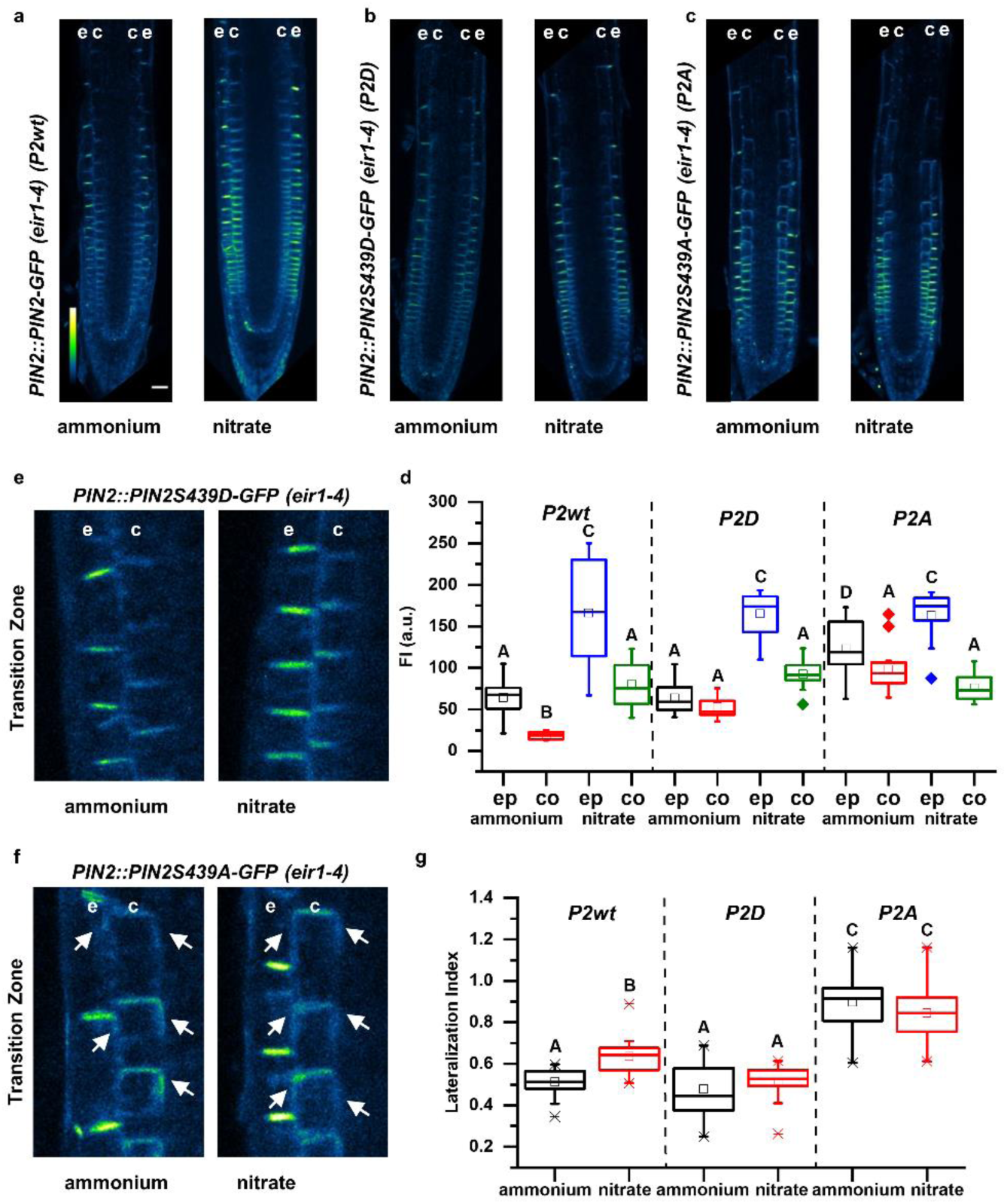
Impact of Ser439 on PIN2 localization in roots supplemented with ammonium or nitrate. **a - c**. Pseudo-colored, optical longitudinal cross sections of 5 DAG roots expressing (a) PIN2-GFP (*PIN2::PIN2-GFP, P2wt*) (**b**) *PIN2S439D-GFP* (*PIN2::PIN2S439D-GFP, P2D*) and (**c**) *PIN2S439A* (*PIN2::PIN2S439A-GFP, P2A*) - all in *eir1-4* background – 12 HAT to ammonium or nitrate supplemented media. “e” denotes epidermis and “c” cortex, respectively. Color code represents GFP intensity from low (blue) to high (white) values. Scale bar = 50 µm. **d**. Box plots display the distribution of the cell membrane derived PIN2-GFP fluorescence intensity (FI) values (in arbitrary units, a.u.) in roots transferred to ammonium ((grey, epidermis (ep) and red, cortex (co) and to nitrate (blue, epidermis (ep) and green, cortex (co)). 5 cells per roots were analyzed in at least 9 roots per genotype per treatment. The statistical significance was evaluated with ANOVA at p<0.05. **e and f**. Microscopic images of 5 DAG old roots expressing (**e**) *PIN2::PIN2S439D-GFP* and (**f**) *PIN2::PIN2S439A-GFP* 12 HAT to ammonium or nitrate amended media. “e” denotes epidermis and “c” cortex, respectively. White arrows point to PIN2-GFP protein localization on the lateral membranes. **g**. Box plots display lateralization index (fluorescent signal detected on apical/basal membranes vs inner/outer membranes) of *P2wt, P2D* and *P2A* roots transferred to ammonium (grey) or nitrate (red) supplemented medium. At least 24 cells from 5 roots were analyzed per genotype per treatment. The statistical significance was evaluated with ANOVA at p<0.05.

### Nitrogen dependent PIN2 phosphorylation fine-tunes PIN2 mediated root growth

Next, we examined the impact of PIN2 phosphorylation status on the root growth adaptations to different N source. To evaluate functionality of PIN2-GFP constructs with phosphosite substitutions, we analyzed their ability to rescue the agravitropic phenotype of *eir1-4. PIN2::PIN2*^*S439D*^*-GFP* as well as *PIN2::PIN2*^*S439A*^*-GFP* constructs were able to rescue the agravitropic phenotype of the *eir1-4* mutant (Fig. S8a), indicating that the overall activity was maintained in both mutated variants. However, measurements of roots 6, 24 and 96 HAT on either NH_4_^+^ or NO_3_^-^ supplemented media revealed that modulation of PIN2 phosphorylation status interfere with the flexible adjustment of root growth to N source. Roots of *PIN2::PIN2*^*S439D*^*-GFP,eir1-4* exhibited enhanced growth already 6 hours after transfer on NO_3_^-^ when compared to NH_4_^+^ supplemented medium (Fig. 7a); however, when compared to control seedlings the enhancement of root growth by NO_3_^-^ was less pronounced 4 DAT (Fig. S8b). This suggests that PIN2^S439D^-GFP is partially able to mediate distinct root growth responses to different N sources. Roots of *eir1-4* expressing *PIN2::PIN2*^*S439A*^*-GFP* exhibited delay in adjusting growth to NO_3_^-^ provision and no significant increase in length 6 and 24 HAT to NO_3_^-^ when compared to NH_4_^+^ could be detected (Fig. 7a; S8b).

**Figure 7.**
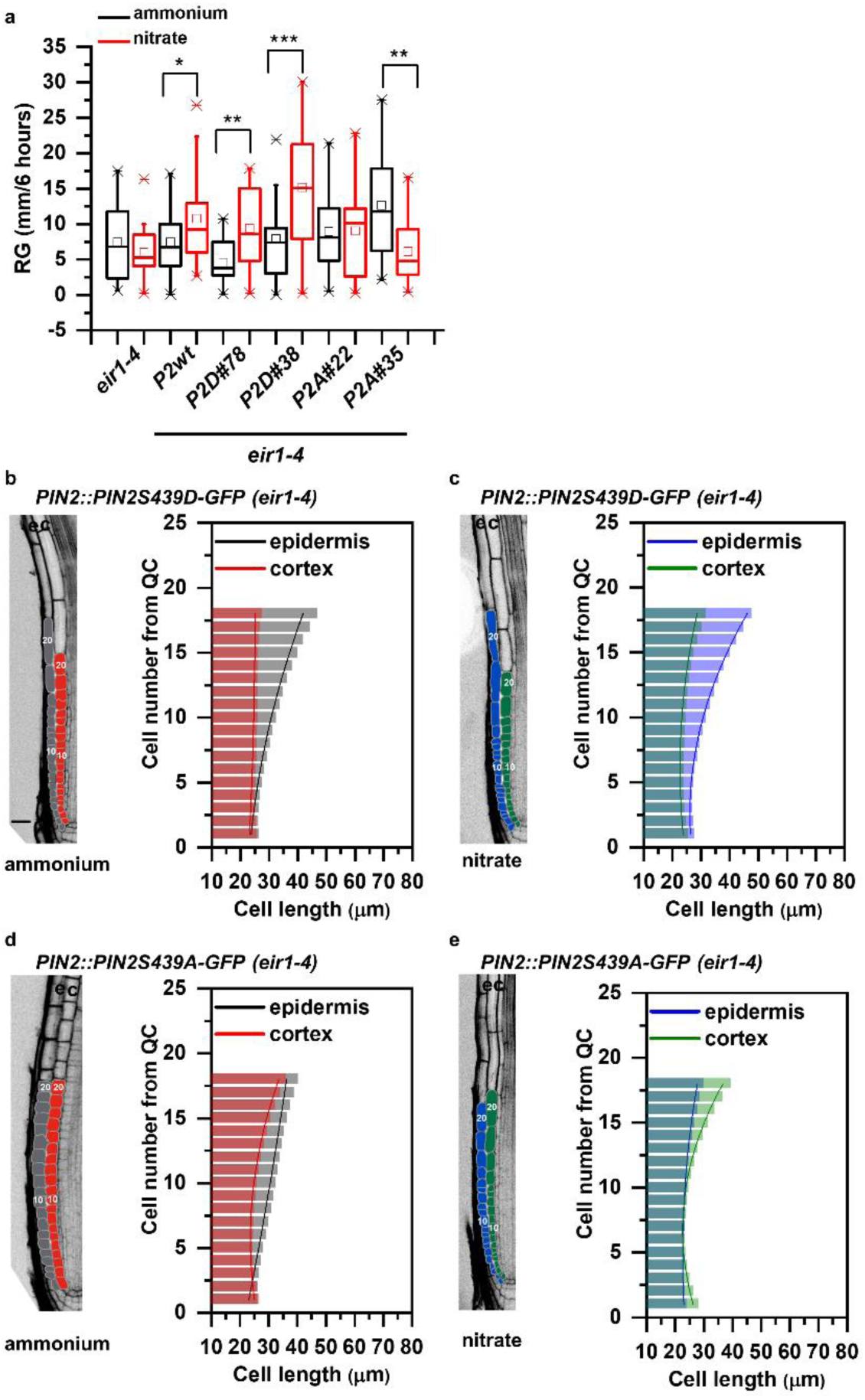
Impact of PIN2S439 phospho-variants on the adaptation of the primary root growth to ammonium or nitrate provision. **a**. Box plot representation of root growth (µm/6 hours) of *eir1-4*, Col-0, *PIN2::PIN2-GFP* (*P2wt*), two independent *PIN2::PIN2S439D-GFP* (*P2D*) lines (#78 and #38) and two independent *PIN2:PIN2S439A-GFP* (*P2A*) lines (#22 and #35) transferred to ammonium or nitrate containing medium. At least 10 roots were analyzed per genotype per treatment. The statistical significance was evaluated with ANOVA at p<0.05. **b - e**. Optical, longitudinal sections of 5 DAG old roots expressing *PIN2S439D-GFP* (**b and c**) and *PIN2S439A-GFP* (**d and e**) 12 HAT to ammonium (**b and d**) or nitrate (**c and d**) supplemented media. The first 20-20 epidermal and cortex cells (from quiescent center (QC)) are highlighted in grey and in red on ammonium (**b and d**) and in blue and green on nitrate (**c and e**), respectively. Scale bar = 30 µm. Column bars denote the geometric mean of cell length at the respective positions. Lines represent a polynomial regression fit, with calculated slopes between cells 10 and 20 of ammonium-PIN2S439D-epidermis: 1.38867+0.03079, ammonium-PIN2S439D-cortex: 0.05689+0.00497, nitrate-PIN2S439D-epidermis: 1.92749+0.0727, nitrate-PIN2S439D-cortex: 0.66477+0.03592, ammonium-PIN2S439A-epidermis: 0.7164±0.00565, ammonium-PIN2S439A-cortex: 1.09064±0.05609, nitrate-PIN2S439A-epidermis: 0.53796±0.0249, nitrate-PIN2S439A-cortex: 1.61118±0.09541. Data are derived from 3 independent experiments; at least 5 roots were analyzed in each case.

Intriguingly, although *PIN2::PIN2*^*S439D*^*-GFP* partially recovered the ability of *eir1-4* roots to adjust elongation growth to N source, in depth analysis of epidermal and cortex cell files revealed intriguing differences when compared to control roots. In *PIN2::PIN2*^*S439D*^*-GFP;eir1-4* roots, irrespective of the N source, length of epidermal cells steeply increased with distance from QC, whereas cortex cells underwent slow steady elongation. Similar asynchronous growth patterns in epidermal and cortex cell files were observed in NH_4_^+^, but not in NO_3_^-^ supplied *PIN2::PIN2-GFP;eir1-4* and Col-0 roots, indicating that *PIN2::PIN2*^*S439D*^*-GFP* is not able to recover all aspects of root adaptation to varying N supply (Fig. 7b,c compared to Fig. S8c, Fig.1c). In *PIN2::PIN2*^*S439A*^*-GFP* roots irrespective of the N source, shallow slope of epidermal cell length was detected, which resulted in synchronized growth patterns of epidermal and cortex cell files, resembling those observed in *PIN2::PIN2-GFP,eir1-4* and Col-0 roots supplied with NO_3_^-^ (Fig. 7d, e compared to Fig. S8c, Fig.1d). Thus, *PIN2::PIN2*^*S439A*^*-GFP, eir1-4* roots supplemented with NH_4_^+^ acquired features typical for Col-0 roots supplied with NO_3_^-^.

In summary, the PIN2^S439D^-GFP phospho-variant is lacking the enhanced elongation growth of *eir1-4* roots, but it is unable to synchronize the patterns of epidermis and cortex elongation in response to NO_3_^-^. Unlike PIN2^S439D^-GFP, PIN2^S439A^-GFP is unable to rescue sensitivity of *eir1-4* roots to NO_3_^-^ stimulatory effect on root elongation growth and to synchronize the patterns of epidermis and cortex elongation growth, irrespective of N source. Taking together, these results indicate that N dependent regulation of phosphorylation status of PIN2 at S439 is a part of complex mechanism underlying root growth adaptation to specific N source, which involves coordination of tissue specific balance between cell proliferation and elongation.

### An experimentally-derived quantitative model predicts nitrogen-dependent coordination of root growth

Experimental findings suggest that nitrogen-dependent fine-tuning of polar auxin transport through the regulation of PIN2 phosphorylation status could coordinate the growth of adjacent tissues and thereby steer the root growth. To mechanistically understand nutrient effect on plant root growth, we developed a multilevel computer model of epidermis and cortex tissues. The complete scheme of the model components can be found in Fig. S9 and a full description of the model is provided in the Methods section. The model integrates the experimental observations of N source dependent effects on PIN2 accumulation at the PM (Supplementary dataset 1) and previously shown auxin-dependent degradation of PIN2^51,52^. As a source of auxin we tested two likely scenarios *i.)* a uniform source of auxin along the epidermis (Model A) or *ii.)* flow of auxin from lateral root cap (LRC) and QC into epidermis^53,54^ (Model B). In addition, other less favorable scenarios were also tested (see Methods and Fig. S10f). Importantly, PIN2 polarity and auxin distribution as well as cell length and number of cell division resulted purely from predictions of the model. To test our models, we compared experimental observations of PIN2 distributions (Supplementary Video 6, Supplementary data set 1), cell length measurements (Fig. 1, Supplementary dataset 1) and auxin content (Fig. 2, Supplementary data set 1) with the predicted by computer model simulations. The initial Model A failed to recapitulate experimental data (Fig. S10b-d), indicating that the auxin source assumption may not be correct and/or there are missing components, which were not considered in this model. Model B, which unlike Model A, integrates flow of auxin from the QC and the LRC into the epidermis, and in addition, it implements correlation between cell distance from the QC with both increased PIN2 trafficking and PIN2 degradation (Fig. S10e), was able to recapitulate PIN2 and auxin distributions as well as cell length across the meristem (Supplementary Fig.10a-d and 10f). To comprehend a necessity for these two essential components in our model, we closely inspected the relation between auxin activity levels and PIN2 fluorescence in our experimental dataset in roots supplemented with NO_3_^-^ or NH_4_^+^. Our analysis revealed that for the same auxin activity two different PIN2 levels were observed in both the cortex and the epidermis that was dependent on distance from the QC - a component missing in Model A (Fig. S10a). This eminently bi-stable feature was important to guarantee the synchrony of cell elongation between adjacent tissues as this feature was compromised in NH_4_^+^ grown roots that showed an asynchronous elongation of adjacent cortex and epidermal cells (Fig. S10a). Notably, Model B could successfully capture this relation (Fig. S10c, d). Finally, we coupled auxin activity to cell division and elongation and simulated our root model in both NH_4_^+^ and NO_3_^-^ regimes (Fig. 8b). As for previous simulations, the computer model of root growth does not include neither fixed auxin levels nor pre-patterned PIN2 polarization and was capable of recapitulating *in planta* root growth patterns in those different N sources (Fig. 8b, Supplementary Videos 7 and 8). Furthermore, model predictions such as lateral auxin distribution (Fig. 8c), meristem length (Fig. 8d) and proliferation dynamics (Fig. 8e) are in a fair agreement with experimental results (Figs. 1 and 2). Importantly, our model predicts mechanistic principles of the growth in synchrony such as coordinated cell divisions in both epidermis and cortex tissues further away from QC (Fig. 8e) and lateral auxin transport through PIN2 between cortex and epidermis near the transition zone (Fig. 8c).

**Figure 8.**
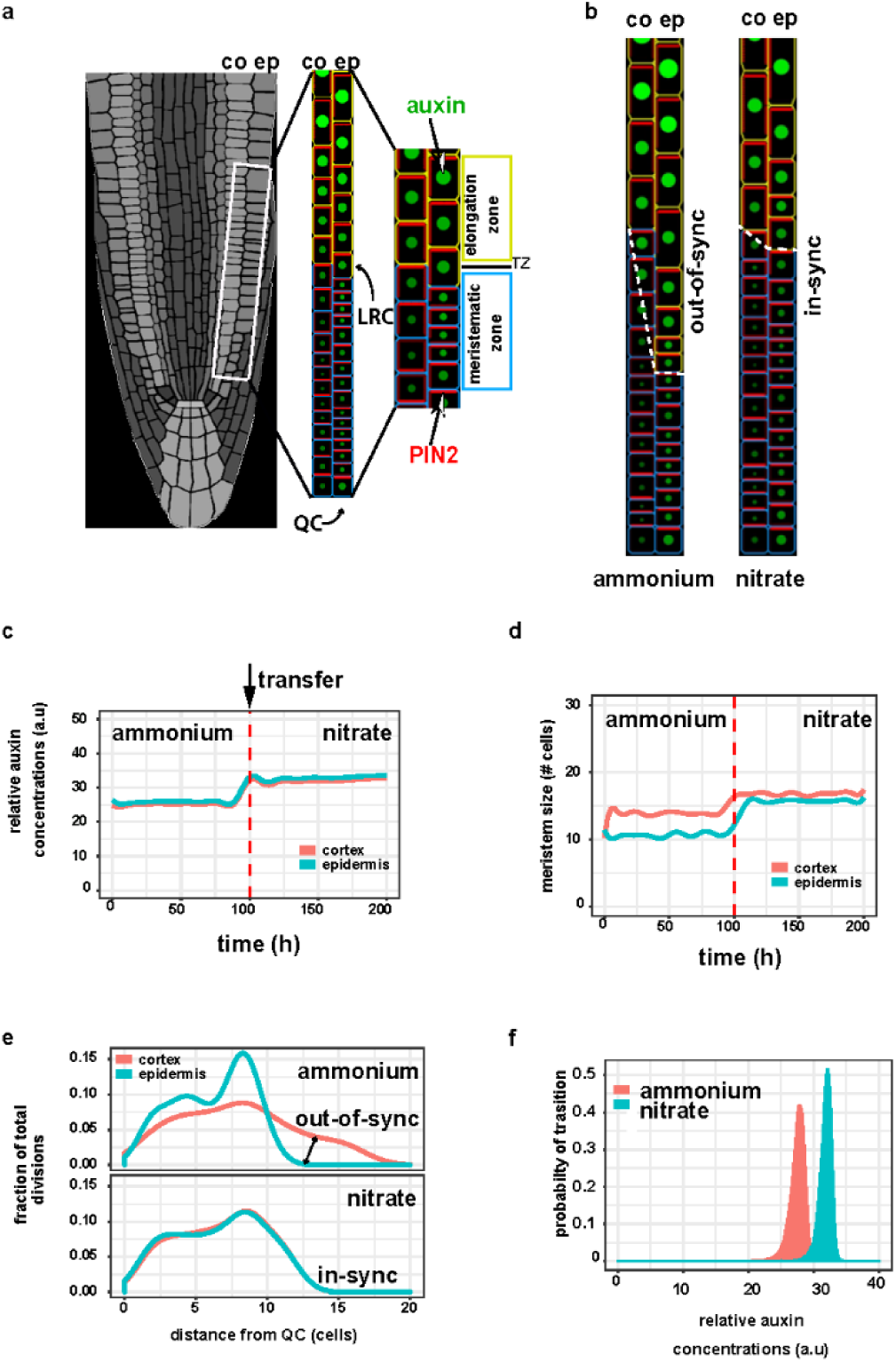
Dynamic computer model of root growth predicts nitrogen source dependent effect on cell growth dynamics, auxin distribution and root zonation. **a**. Schematics of the root model with epidermis (ep) and cortex (co) tissues. Meristematic and elongating cells are shown with blue and yellow walls, respectively. Auxin levels are represented by green circle size and red bars reflect the PIN2 amounts. Auxin is supplied from Lateral Root Cap (LRC) and QC (Model B) **b**. Steady state snapshots from model simulation with ammonium (left panel) and with nitrate (right panel). Note out-of-sync growth patterns (dashed white line) in ammonium. **c and d**. Model simulation representing the effect of the transition from ammonium to nitrate (denoted by a red dashed line) on the relative level of auxin (C) and meristem size measured as distance from QC (D). **e**. Model predictions display the fraction of total cell division events per cell in the meristem along in the two N source. Note cell division is out-of-sync in ammonium, producing altered growth of the root. **f**. Experimentally derived relative auxin level threshold triggering cell elongation depends on the actual N content of the root.

Mechanisms that trigger the transition from meristematic activity to cell elongation are not well understood^55^. Auxin plays a fundamental role in the establishment of the TZ^56,57^. Our model could predict a precise threshold of auxin levels that was necessary to determine the transition to elongation. This auxin threshold is dynamic as it depends on the actual N source; in particular, higher levels of auxin were required to advance cell elongation on NO_3_^-^ (Fig. 8f). Taken together, we have developed a quantitative experimentally-supported computer model of root growth in different N sources that was capable of recapitulating all experimental observations as well as generating new predictions that could broaden our understanding of root growth mechanisms in the dynamic environment.

## Discussion

Ammonium and nitrate represent major inorganic forms of N absorbed by plants. Since the distribution of these N sources in the soil is very heterogeneous^58^, plants tend to maximize the N exploitation by flexible modulation of root system architecture^59^. Although distinct impacts of NH_4_^+^ and NO_3_^-^ on the root system growth and development have been already demonstrated^6^, molecular mechanisms how spatio-temporal changes in N resource impact on root growth are scarcely described.

Root growth is determined by the production of new cells at the apical meristem and their rapid elongation need to be well coordinated across diverse cell types of the root organ. Conversion from proliferative to elongation phases occurs as cells pass through the TZ where they undergo complex cyto-architectural re-arrangement^57^. Hence, alterations of root growth kinetics might result from modulation of any of these growth-determining processes. We show that replacement of NH_4_^+^ for NO_3_^-^ has a rapid impact on root growth kinetics and in particular progression of cells through individual root zones. While in roots supplied with NH_4_^+^, proliferative capacity of epidermal cells is attenuated in closer distance to the QC, which led to their earlier and rapid transition to elongation phase when compared to cortex, provision of NO_3_^-^ promotes proliferation and steady elongation of epidermal cells, which results in well-synchronized growth patterns of epidermis and cortex. Hence, adaptation of primary roots to different sources of N encompass a tissue specific modulation of cell proliferation and cell growth kinetics.

Auxin is an essential patterning cue during plant growth and development. A number of recent studies have demonstrated that levels and distribution of this hormone have instructive function in many aspects of root growth including the root apical meristem patterning, its size determination, transition of meristematic cells into the elongation phase and capacity of cells to elongate^55,60,61^. Whereas the exit of cells from meristematic zone was associated with local auxin minima that has been proposed to define the transition zone^60^, increase of auxin signaling along the longitudinal root growth axis correlated with cell wall acidification as a potential driving force of cell elongation^62^.

Experimental measurements supported by a quantitative computational model indicate that adjustment of root growth dynamics in different N regimes is dependent on the precise modulation of auxin transport routes between cortex and epidermis. The steep increase of auxin activity correlating with earlier attenuation of proliferation activity in the epidermis and transition of cells into the elongation phase was eminent in roots grown on NH_4_^+^. In contrast, shallow slopes of the auxin activity in both epidermis and cortex corresponded with delayed, gradual transition of epidermal cells into elongation phase in roots supplemented with NO_3_^-^, showing a tight growth synchronization with adjacent cortex tissues. Based on these observations we demonstrate that a flexible modulation of auxin activity in response to varying sources of N is largely consistent with described impact of auxin on key events defining root growth such as transition into elongation growth and kinetics of elongation.

Delivery of auxin in outer tissues including the cortex and the epidermis is largely mediated by the PIN2 auxin efflux carrier^43,44^. While PIN2 dependent basipetal transport of auxin is instructive for elongation growth and root gravity bending^63^, PIN2 mediated reflux to inner tissues has been associated with maintenance of root meristem size^39^. Measurements of the auxin transport revealed that replacement of NH_4_^+^ by NO_3_^-^, significantly enhances flow of auxin in the basipetal direction which correlates with increased PIN2 activity near the transition zone. Loss of PIN2 activity not only interferes with the NO_3_^-^ stimulated transport of auxin towards the shoot, but also severely affects adaptive responses of roots to this N source. Furthermore, model predictions based on these experimental measurements suggest a bi-stable relationship between auxin levels and PIN2 activity and cell elongation that is enhanced in NO_3_^-^, which could explain why roots grown on nitrate can coordinate their growth by passing auxin between cortex and epidermal cells in a synchronous manner. Furthermore, our model confirmed the necessity for self-emerging communication between cortex and epidermis via auxin with quantitative computer simulations of root growth under different N conditions.

Dynamic, N source dependent accumulation and polarization of PIN2 at the PM, but unchanged *PIN2* transcription, pointed at post-transcriptional regulatory mechanism underlying adaptation of basipetal auxin transport to N supply. Replacement of NH_4_^+^ by NO_3_^-^ promoted accumulation of PIN2 at the apical PM of epidermal and cortex cells as well as to the lateral sides.

Phosphorylation has been recognized as a prominent posttranslational modification of PIN proteins that determines their polar membrane localization and activity^64^. Unexpectedly, genome-wide analysis of the phosphoproteome during early phases of root adaptation to provision of NO_3_^-^ (Vega et al.) retrieved PIN2 among differentially phosphorylated proteins. Serine 349 of PIN2 in *Arabidopsis*, found to undergo a rapid de-phosphorylation after replacement of NH_4_^+^ by NO_3_^-^. The PIN2S439 phosphosite was not completely unknown: it was originally identified as differentially phosphorylated during lateral root morphogenesis^65^. It is positioned in the hydrophilic loop domain of the PIN2 protein and is an evolutionarily conserved residue in the PIN2 or PIN2-like clade across species including gymnosperms, mono- and dicotyledonous plants, suggesting that PIN2 might be universally involved in other plant species adaption strategies to the changing N sources by means of its post-translational (phosphorylation) mechanism. The functional characterization of PIN2 and its phosphor-variants suggests that N source dependent regulation of PIN2 phosphorylation status has a direct impact on the flexible adjustment of PIN2 membrane localization and polarity, and thereby adaptation of root growth to varying forms of N supply.

## Acknowledgments

We acknowledge Gergely Molnár for critical reading of the manuscript, Alexander Johnson for language editing and Yulija Salanenka for technical assistance. Work in the Benková lab was supported by the Austrian Science Fund (FWF01_I1774S) to KÖ and EB. Work in the Wabnik lab was supported by the Programa de Atracción de Talento 2017 (Comunidad de Madrid, 2017-T1/BIO-5654 to K.W.), Severo Ochoa Programme for Centres of Excellence in R&D from the Agencia Estatal de Investigación of Spain (grant SEV-2016-0672 (2017-2021) to K.W. via the CBGP) and Programa Estatal de Generación del Conocimiento y Fortalecimiento Científico y Tecnológico del Sistema de I+D+I 2019 (PGC2018-093387-A-I00) from MICIU (to K.W.)

We acknowledge the Bioimaging Facility in IST-Austria and the Advanced Microscopy Facility of the Vienna BioCenter Core Facilities, member of the Vienna BioCenter Austria, for use of the OMX v4 3D SIM microscope. AJ was supported by the Austrian Science Fund (FWF): I03630 to J.F.

## Author Contributions

K.Ö. and E.B. conceived the project; K.Ö. performed most of the experiments; M.M. contributed to the generation of the computational model; A.V., J.B. and R.G. shared unpublished material; A.J. performed the 3D-SIM experiment; R.A. performed q-PCR and GUS-staining experiments; L.A. performed the regression analysis; J.C.M. contributed to the multiphoton microscopy imaging; Y.Z. conducted protein sequence alignments; S.T. generated the DII and mDII lines in the *eir1-4* background; C.C., El.B. and A.G. designed and performed some of the pre-pilot experiments; C.A. assisted K.Ö. in multiple experiments; J.F. financially supported A.J., Y.Z. and S.T. The manuscript was written by K.Ö., K.W. and E.B.

---

**Supplementary Figure 1 related to Figure 1.**
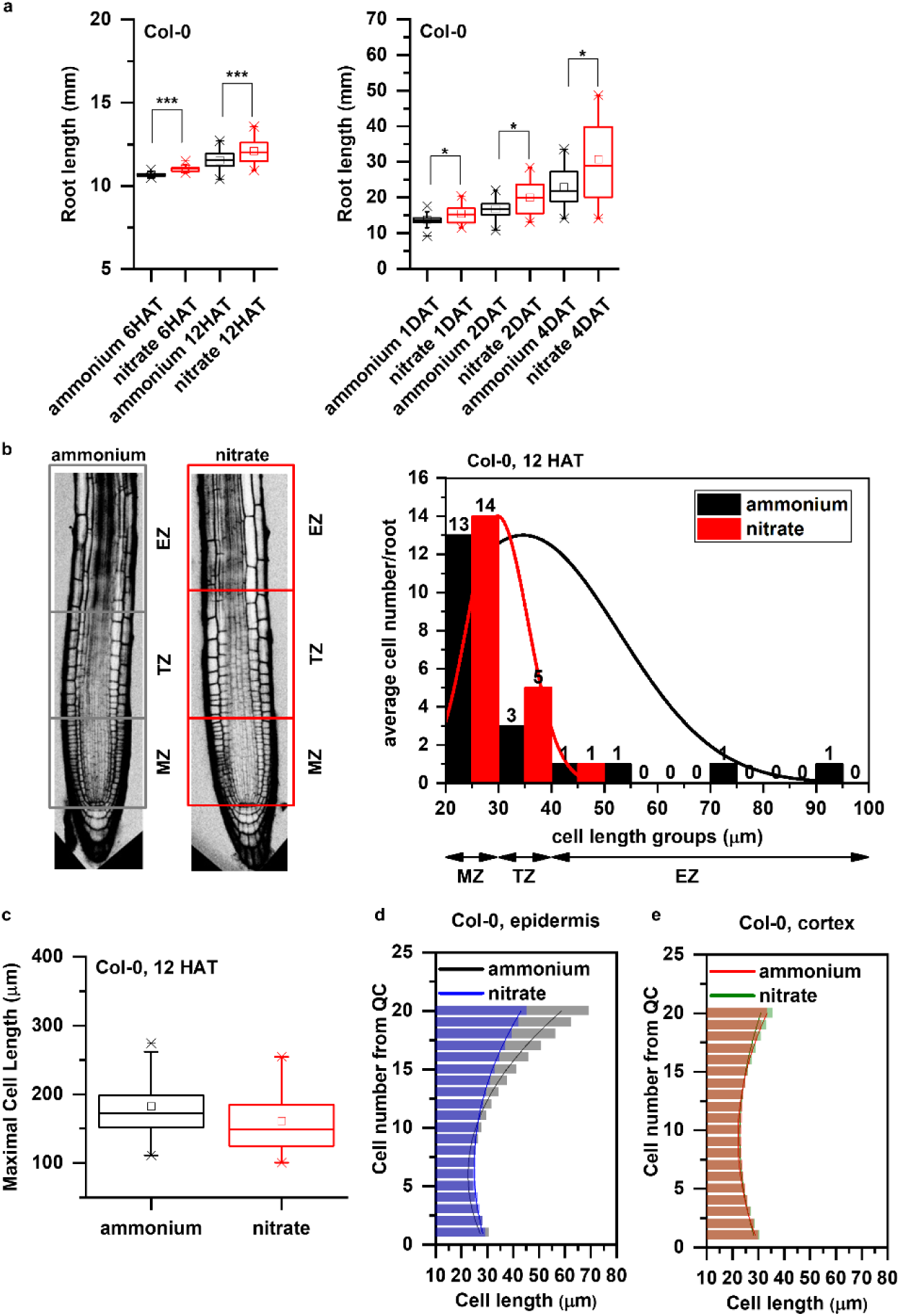
Additional data supporting the distinct growth kinetics of Col-0 roots transferred to ammonium or nitrate supplemented media. **a**. Primary root length (mm) of Col-0 seedlings 6 and 12 HAT and 1, 2, 4 days after transfer (DAT) to ammonium (black) or nitrate (red) supplemented medium. At least 34 roots were measured per time point per treatment. The statistical significance was evaluated with ANOVA at p<0.05. **b**. On the left, schematic representation of distinct root zones: Meristematic Zone (MZ), Transition Zone (TZ, which is interpolated between the apical meristem and the subapical elongation zone) and Elongation Zone (EZ). Boxes highlight the borders of the specified root zones (grey box for ammonium and red for nitrate). On the right, Col-0 epidermal cells length was measured along the root tip (from QC), grouped based on their cell length (x axis) and were plotted against the average cell number per group per root (y axis) in both conditions (ammonium, black and nitrate, red). Note the higher cell number in case of nitrate (red) in the group (30-40 µm, representing TZ). Data are derived from 3 independent experiments, total number of analyzed roots are n=18 in each case. **c**. Maximal cell length (measured at the end of the elongation zone) of Col-0 roots 12 HAT either to ammonium (black) or nitrate (red). 13 roots per treatment, 3 cells per root were analyzed. **d and e**. Comparison of cell length measurements along epidermis (**d**) and cortex (**e**) upon ammonium (black and red) and nitrate (blue and green) treatments. Column bars denote the geometric mean of cell length at the respective positions. Lines represent a polynomial regression fit. Data are derived from 3 independent experiments, total number of analyzed roots are n=18 in each case.

**Supplementary Figure 2 related to Figure 1.**
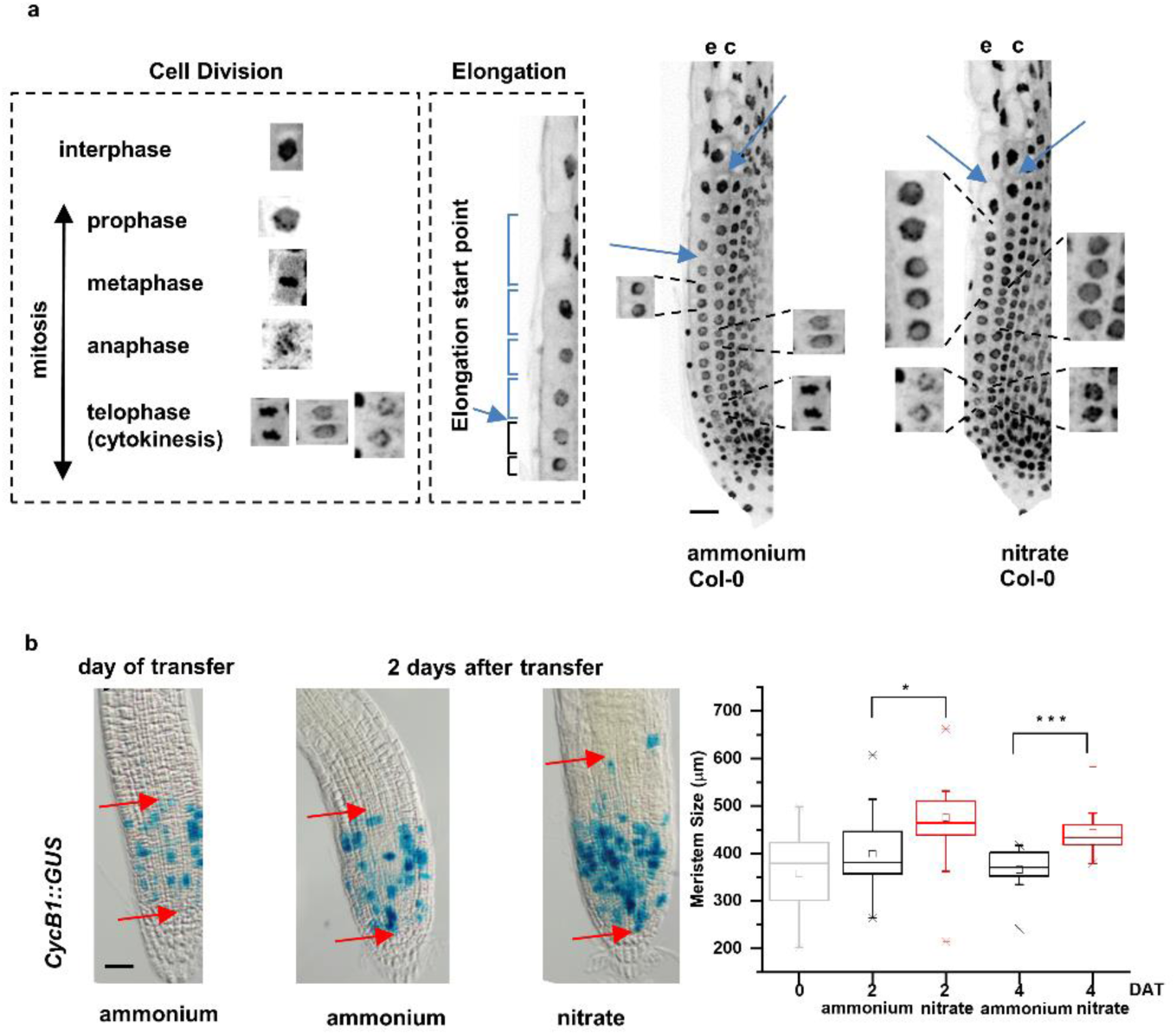
Additional data supporting the distinct division patterns in Col-0 roots on media supplemented with different nitrogen source. **a**. On the left, schematic representation of the different cell division phases of DAPI stained Col-0 roots and an illustration of how the onset of cell elongation was marked. On the right, DAPI stained confocal microscopic images of Col-0 roots 12 HAT to ammonium or nitrate containing medium. Mitotic events are displayed along epidermis (e) and cortex (c). Blue arrows point to the first elongating cells. Scale bar = 50 µm. **b**. Bright field microscopic images of *GUS* expressing roots driven by *CycB1* promoter. Blue spots mark *CycB1* promoter activity. Red arrows point to the beginning and to the end of GUS expressing area (meristem size). Scale bar = 100µm. Box plot chart represents the meristem size (µm) of *CycB1::GUS* expressing roots on the day of transfer (0) and 2 and 4 DAT to ammonium or nitrate. Differences of the means were calculated with a t-test (p value *<0.05, ***<0.001). At least 14 roots were analyzed per time point per treatment.

**Supplementary Figure 3 related to Figure 2.**
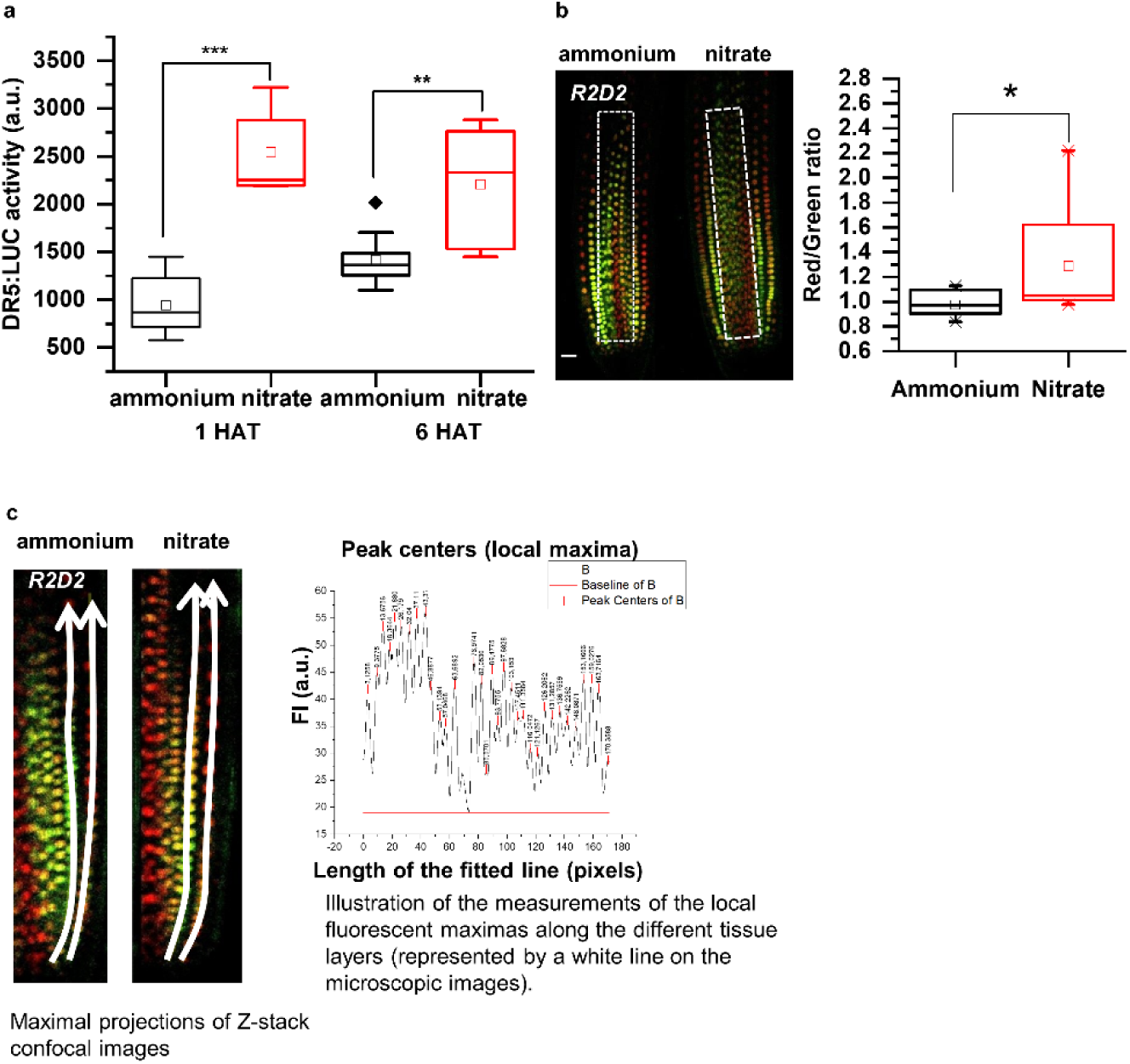
Supporting data for auxin activity and distribution in roots transferred to different nitrogen sources. **a**. *DR5* activity in *Arabidopsis* roots. Box plots represents luciferase activities in *DR5::LUC* expressing roots 1 and 6 HAT. 40 roots were collected per treatment per time points. Experiment was repeated 3 times. Statistical differences were calculated with a t-test (p values **<0.01, ***<0.001) **b**. Expression profile of the auxin-input reporter *R2D2* in the stele (labeled with “white box”) of roots 12 HAT to ammonium or nitrate containing media. Box plots represent the quantification of the red (auxin-independent) vs green (auxin dependent) fluorescent signal ratio in the stele. At least 13 roots were analyzed and statistical difference was calculated with a t-test (p value <0.05). **c**. Illustrations for R2D2 quantification along epidermal and cortical cell files. For details see the “Quantification of R2D2, DII-VENUS and mDII-VENUS fluorescence signal” in the “Methods” section.

**Supplementary Figure 4 related to Figure 3.**
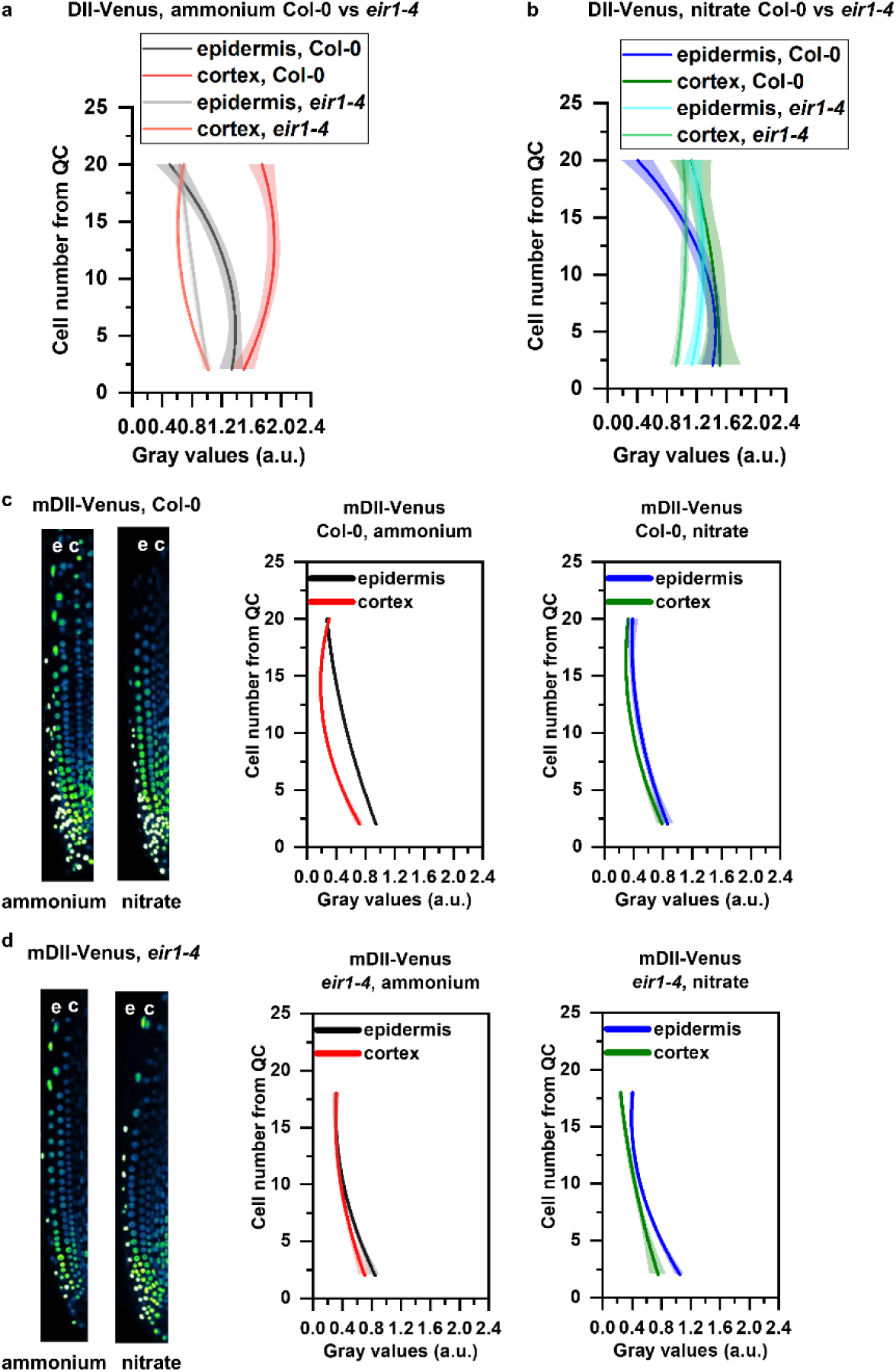
Additional data for demonstrating the distinct patterns of auxin distribution in Col-0 and *eir1-4* roots. **a and b**. Comparison of DII-Venus fluorescent signal in Col-0 and *eir1-4* lines on ammonium (**a**) and on nitrate (**b**) transferred roots. Graphs denote normalized relative auxin levels at the respective positions. Lines represent polynomial regression fit with 95% confidence band. Data are derived from measurements of n=8 (ammonium) and n=10 (nitrate) roots of Col-0 and n=10 roots of *eir1-4* per condition. **c and d**. Maximum intensity Z-stack projection images of 5 DAG old Col-0 (**c**) and *eir1-4* mutant (**d**) roots expressing the non-auxin degradable mDII-Venus reporter grown on ammonium and nitrate supplemented media 12 HAT. “e” and “c” marks epidermis and cortex, respectively. Scale bar = 50 µm. Graphs denote grey values (arbitrary units - a.u.) at the respective positions. Lines represent polynomial regression fit with 95% confidence band. Data are derived from measurements of at least 5 roots per genotype per treatment.

**Supplementary Figure 5 related to Figure 4.**
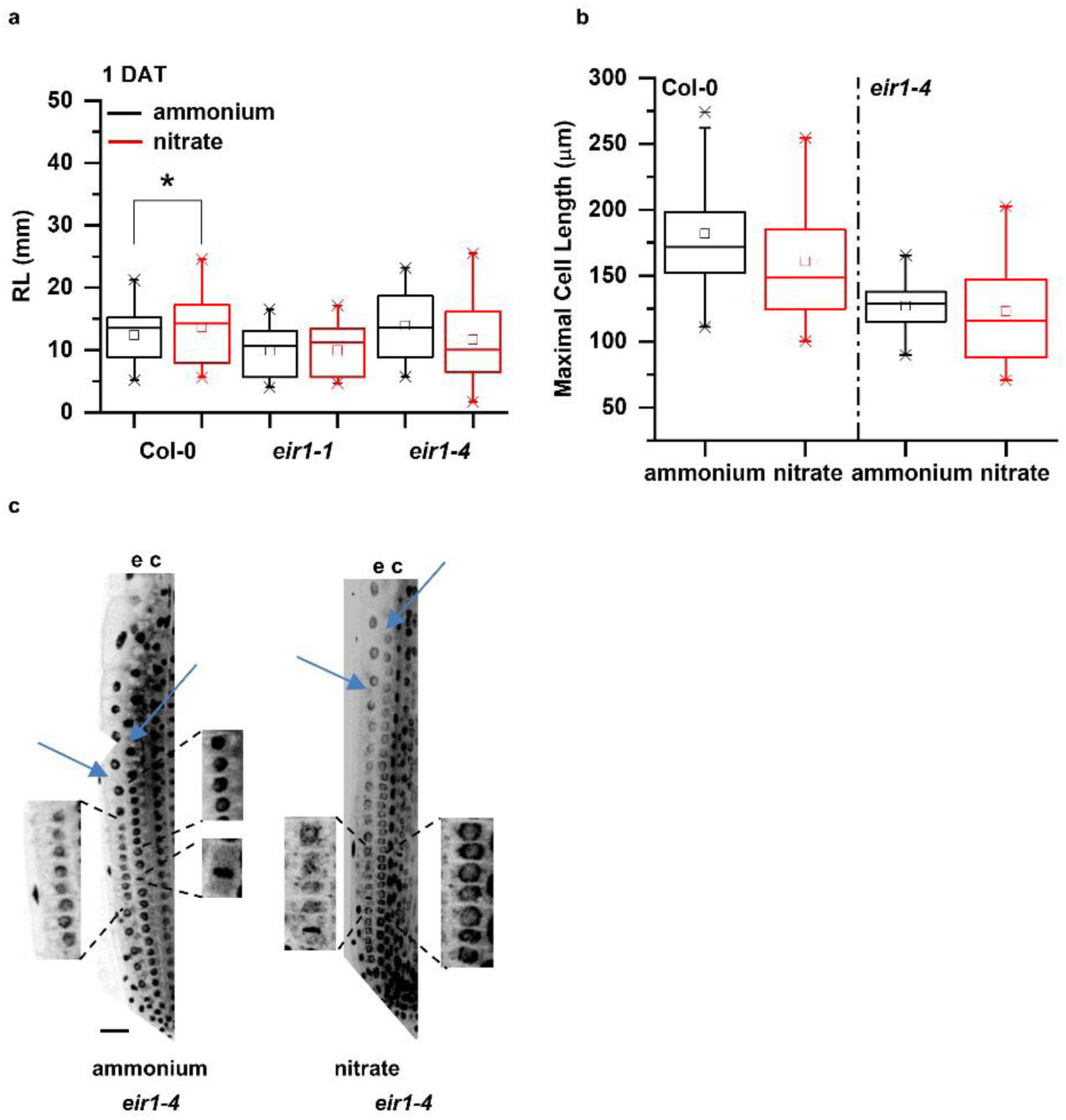
Additional data for demonstrating root growth phenotypes and cell divisions in *pin2* mutants. **a**. Comparison of root length of *pin2* mutants (*eir1-1* and *eir1-4*) to Col-0 on ammonium (black) and nitrate (red) amended media, 1 DAT. At least 11 roots were analyzed per genotype per treatment and statistical difference was calculated with a t-test (p value <0.001) **b**. Box plots of the maximal cell length of Col-0 and *eir1-4* mutant roots 12 HAT to ammonium (black) or nitrate (red). 3-3 cells in at least 13 roots were analyzed per genotype per treatment. **c**. DAPI stained confocal microscopic images of *eir1-4* roots 12 HAT to ammonium or nitrate containing medium. Mitotic events are highlighted along epidermis (e) and cortex (c). Blue arrows point to the first elongating cells. Scale bar = 50 µm.

**Supplementary Figure 6 related to Figure 5.**
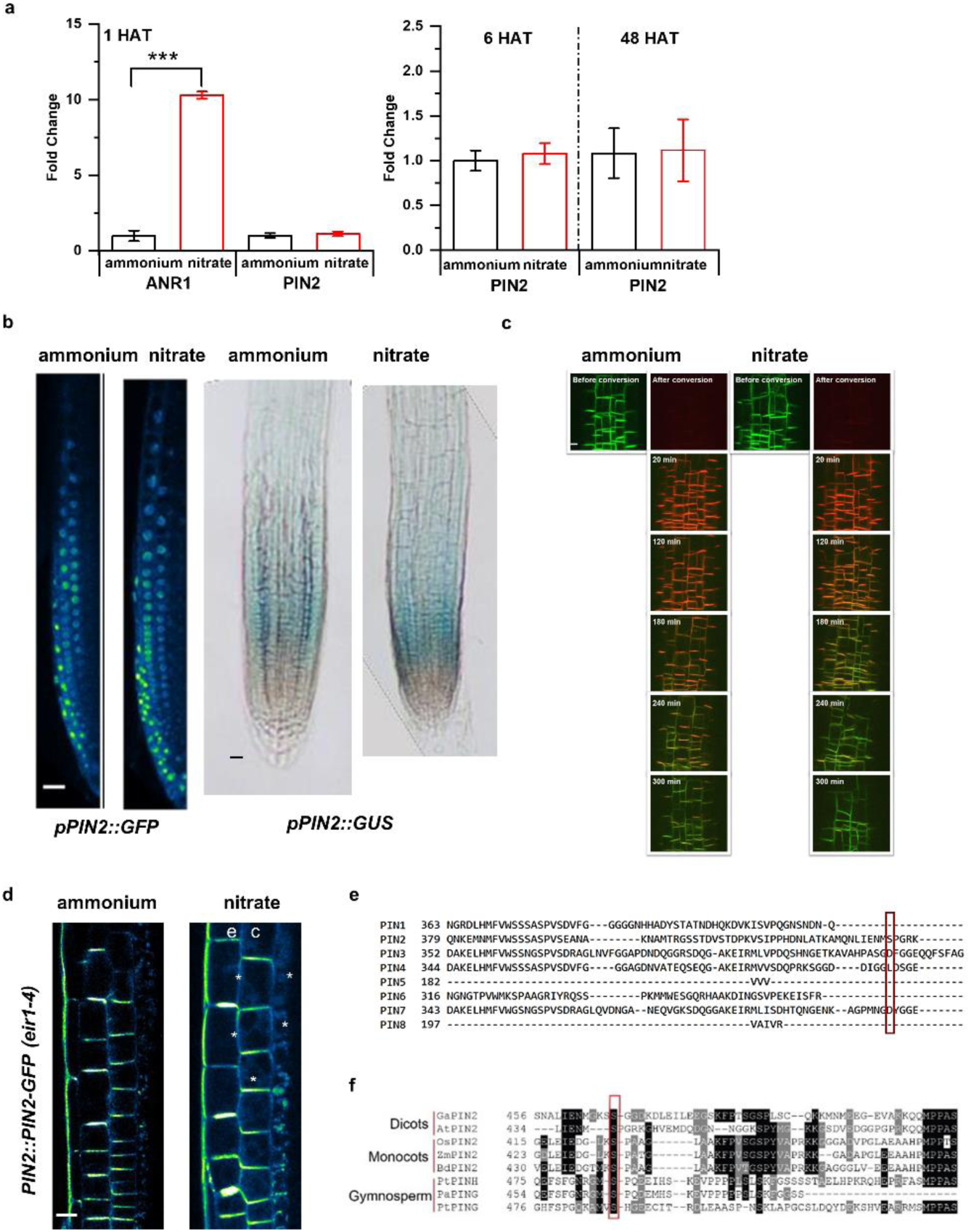
Additional data supporting distinct effects of nitrogen sources on PIN2. **a**. RT-qPCR analysis of *PIN2* expression normalized to *UBQ10 (AT4G05320*) levels in Col-0 roots 1, 6 and 48 HAT to ammonium or nitrate. As a positive control, expression of *ANR1* (nitrate responsive MADS-box transcription factor) was quantified. All RT-qPCR reactions were carried out with biological and technical triplicates. Statistical difference was calculated with a t-test (p value ***<0.001). **b**. *PIN2* promoter activity was monitored in *pPIN2::nlsGFP* and *pPIN2::GUS* expressing roots 12 HAT to ammonium or nitrate. Scale bars = 50 µm. **c**. Confocal microscopic images of *PIN2::PIN2-DENDRA* fluorescence in the same area of the root transition zones 12 HAT to ammonium or nitrate before and after photoconversion (0, 20, 120, 180, 240, 300 min). Scale bar = 20 µm. **d**. Multiphoton microscopic image showing polarity changes of PIN2 expression upon nitrate treatment. “e” and “c” denote epidermis and cortex respectively. White arrows mark lateralization of the PIN2-GFP signal in cortex cells (c). **e**. Protein sequence alignment of members of the *Arabidopsis* PIN protein family. Ser439 of PIN2 and the corresponding residues of other PIN family members are marked by a red box. **f**. PIN2 protein sequence alignment shows evolutionary conservation of Ser439 in representative members of Gymnopserms, Monocots and Dicots. From Gymnosperms *Picea abies* (Pa) and *Pinus taeda* (Pt) PIN2 proteins (PtPING, PtPINH, PaPING), from Monocots *Zea mays* (Zm), *Brachypodium distachyon* (Bd) and *Oryza sativa* (Os) PIN2 proteins (ZmPIN2, BdPIN2, OsPIN2) and from Dicots *Gossypium arboreun* (Ga) and *Arabidopsis thaliana* (At) PIN2 proteins (GaPIN2, AtPIN2) were used. Protein alignments were created with the MEGAX software^66^.

**Supplementary Figure 7 related to Figure 5a.**
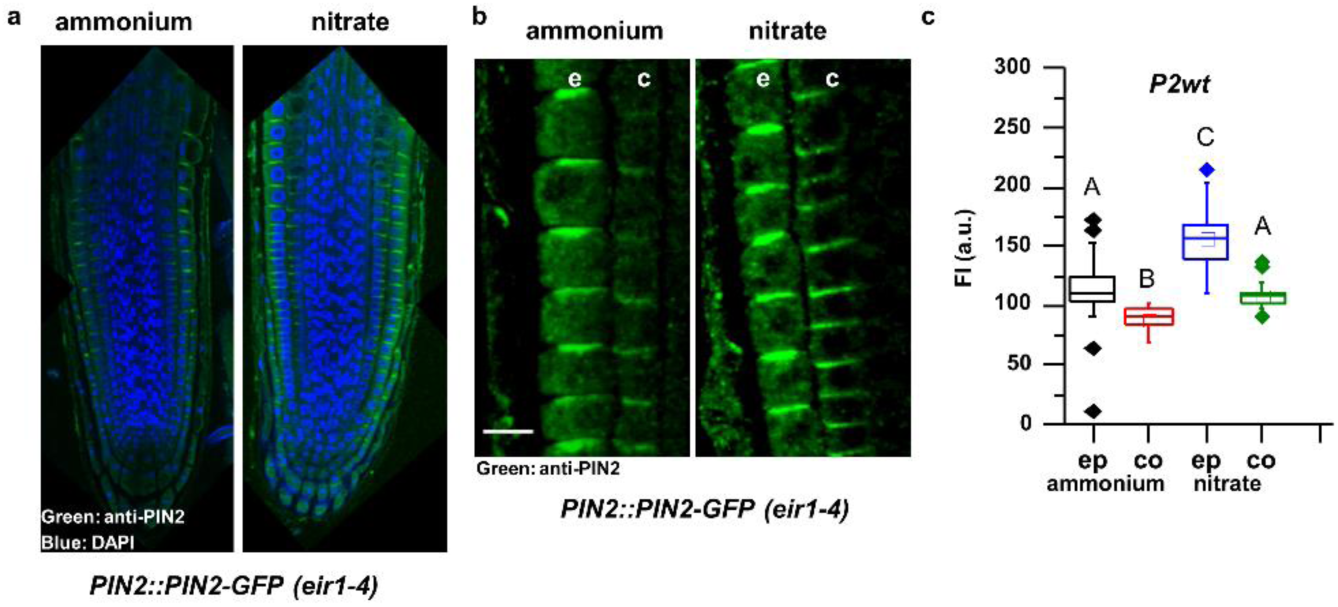
PIN2 immunostaining in *PIN2-GFP* expressing roots. **a**. Confocal microscopic images show anti-PIN2 immunostained PIN2-GFP expressing root tips 12 HAT to ammonium or nitrate. Green and blue signal represents PIN2 and nuclear staining with DAPI, respectively. **b**. Higher magnification of anti-PIN2 immunostained *PIN2-GFP* expressing root in the transition zone. “e” and “c” denote epidermis and cortex, respectively. Scale bar = 25 µm. **c**. Quantification of PIN2 fluorescent signal in the immunostained, ammonium or nitrate treated roots in epidermal (ep) and cortical (co) cell files. Statistical difference was evaluated with ANOVA at p<0.05. 10 roots were analyzed per treatment.

**Supplementary Figure 8 related to Figure 7.**
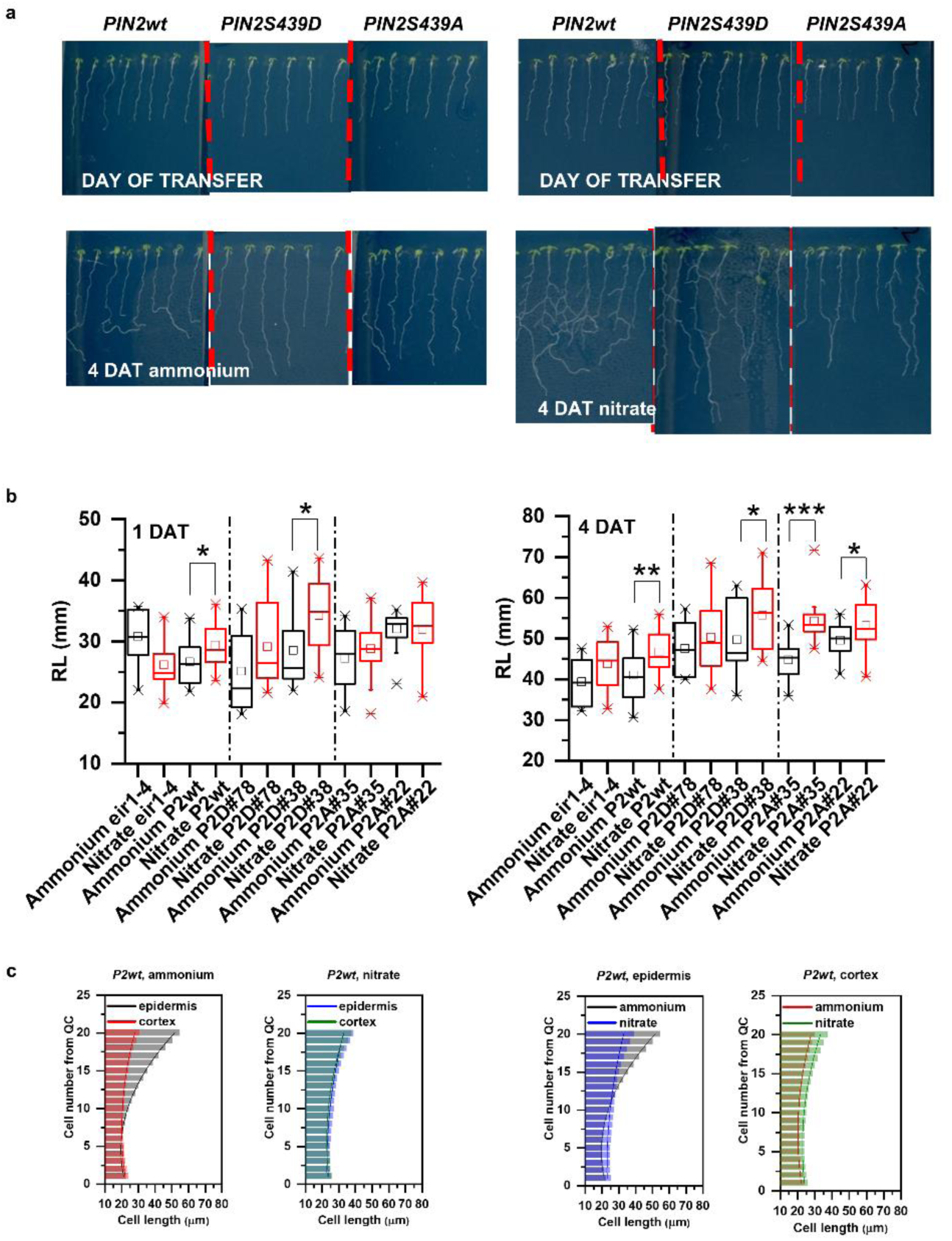
Additional data supporting impact of PIN2S439 phospho-variants on root growth adaptation to source of nitrogen. **a, b**. Seedlings expressing *PIN2::PIN2-GFP* (*P2wt*), *PIN2::PIN2S439D-GFP* (*P2D*) and *PIN2::PIN2S439A-GFP (P2A)* in *eir1-4* background. Representative images of seedlings at the day of transfer and 4 DAT to ammonium or nitrate supplemented plates are shown (**a**). Quantification of root length (mm) in *eir1-4, PIN2wt (eir1-4)* and two independent *P2D (eir1-4)* (#78 and #38) and *P2A (eir1-4)* (#35 and #22) lines 1 and 4 DAT to ammonium or nitrate containing media. At least 7 roots per genotype per treatment were analyzed. Statistical difference was evaluated with a t-test (p values *<0.05, **<0.01 and ***<0.001). **c**. Comparison of cell length changes in epidermal and cortical cell files of *PIN2::PIN2-GFP* (*P2wt*) expressing roots transferred to ammonium or nitrate. Column bars denote the geometric mean of cell length at the respective positions. Lines represent a polynomial regression fit. Data are derived from measurements of 20 roots per genotype per treatment.

**Supplementary Figure 9.**
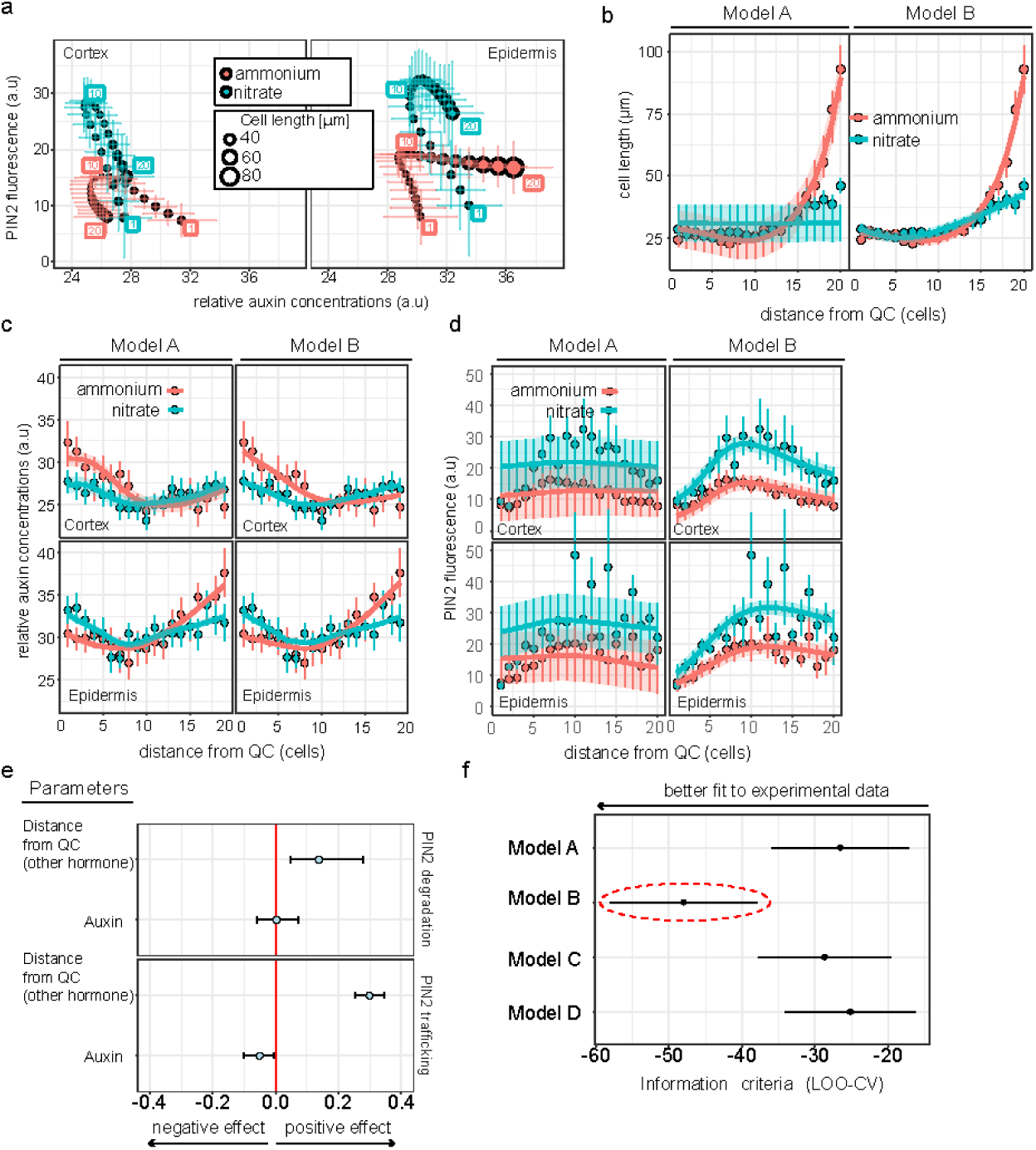
Model robustness and fitting to experimental measurements. **a**. The experimentally-driven PIN2 levels in the function of relative auxin levels for cortex (left panel) and epidermis (right panel), respectively. Relative auxin levels were calculated as follows: log(G/(G+R)) / -0.025, where G and R represent DII-Venus (green) and mDII-Tomato (red) fluorescent signals, respectively. Cell length is denoted by dot size, and the distance of the cell from the QC is labelled with consecutive numbers. Vertical and horizontal lines represent the standard error of measurements. **b-d**. Validations in two regimes demonstrate that Model B faithfully recapitulates all experimental measurements. In these plots model predictions (thick lines and shaded areas for the posterior average and 95% confidence intervals, respectively) are plotted against experimental data (dots and vertical bars for data mean and standard deviations, respectively). **e**. Parameter estimations for Model B suggest antithetic cumulative effects of auxin level and distance from QC on PIN2 dynamics. f. Predictive power of four models for auxin input scenario (A-D). The expected log probability density is used as information criterion. Lower information criterion indicates better posterior predictive performance, and therefore a better fit to the experiments (dashed red ellipse).

**Supplementary Figure 10.**
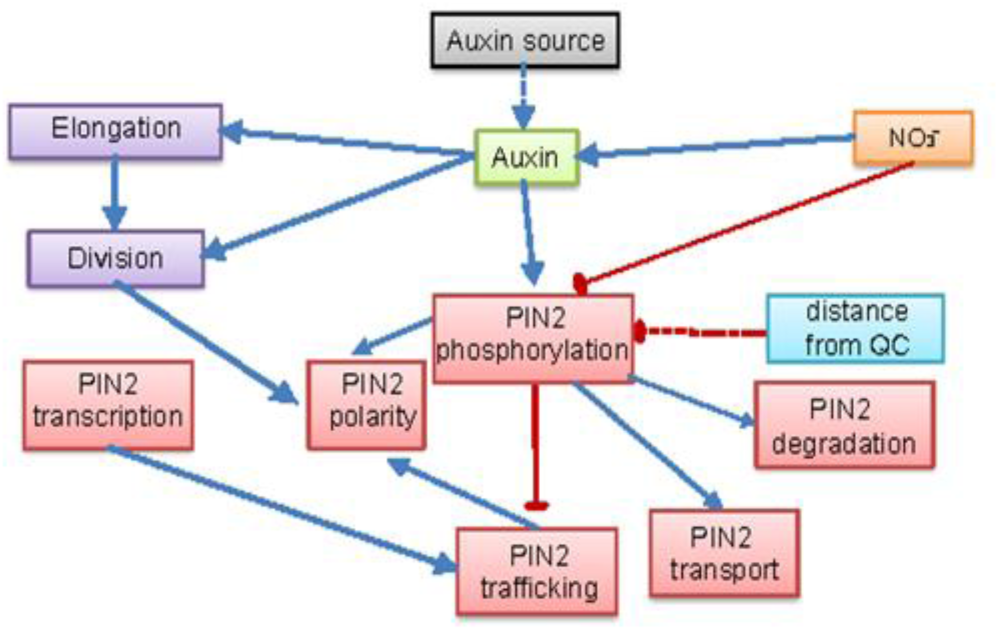
Model diagram. The graphical chart representing the relationships between molecular and structural root processed assumed in the computational model. Solid lines indicated correlation whose effects have been demonstrated (either in our experiments or previous studies). Dashed lines are for relationships that have been integrated in the current model. The two main regulators considered in the model are auxin and nitrate levels, which exert antithetic effect on PIN2 dynamics through phosphorylation.

## Link to Supplementary videos

https://drive.google.com/drive/folders/1AAqNEYuGH2jSOvb_ffvLYYyxyF_eLJSK?usp=sharing

**Supplementary Video 1 related to Figure 1a**.

Time lapse of 5 days old *Arabidopsis* roots expressing the PM marker (*WAVE131Y*) transferred to either on ammonium or nitrate supplemented media and imaged with a vertically oriented LSM700 microscope. Observation of roots initiated 20 minutes after transfer and images recorded every 20 minutes (9 stacks/root/ recording).

**Supplementary Video 2 related to Figure 1b**.

Time lapse of the transition zone of 5-day-old *Arabidopsis* roots expressing the PM marker (*WAVE131Y*) transferred to either on ammonium or nitrate amended media and imaged with a vertically oriented LSM700 microscope. Observation of roots initiated 20 minutes after transfer and images recorded every 20 minutes (9 stacks/root/ recording).

**Supplementary Video 3 related to Figure 5**.

Z-stacks of 2-2 cells in the transition zone of 5-day-old *Arabidopsis* roots expressing *PIN2::PIN2-GFP* 12 HAT to either to ammonium or nitrate amended media and imaged with an Airyscan LSM800 microscope.

**Supplementary Video 4 related to Figure 7**.

Time lapse of *Arabidopsis* seedlings expressing *PIN2::PIN2-GFP* (*PIN2wt*), *PIN2::PIN2S439D-GFP* (*PIN2S439D*) and *PIN2::PIN2S439A-GFP* (*PIN2S439A*). Seedlings were grown on ammonium plates for 7 days and were transferred to ammonium containing agar plates. Plates were scanned on a daily basis with an Epson Perfection V700 flatbed scanner. Images were concatenated with Fiji.

**Supplementary Video 5 related to Figure 7**.

Time lapse of Arabidopsis seedlings expressing wild type *PIN2::PIN2-GFP* (*PIN2wt*), *PIN2::PIN2S439D-GFP* (*PIN2S439D*) and *PIN2::PIN2S439A-GFP* (*PIN2S439A*). Seedlings were grown on ammonium plates for 7 days and were transferred to nitrate containing agar plates. Plates were scanned on a daily basis with an Epson Perfection V700 flatbed scanner. Images were concatenated with Fiji.

**Supplementary Video 6 related to Figure 8**.

A simulation of static example model shows intercellular auxin transport via PIN2 auxin carrier.

**Supplementary Video 7 related to Figure 8**.

Simulation of asynchronous growth of epidermis-cortex tissues in ammonium condition. Related to Fig. 8b.

**Supplementary Video 8 related to Figure 8**.

Simulation of asynchronous growth of epidermis-cortex tissues in nitrate condition. Related to Fig. 8b.

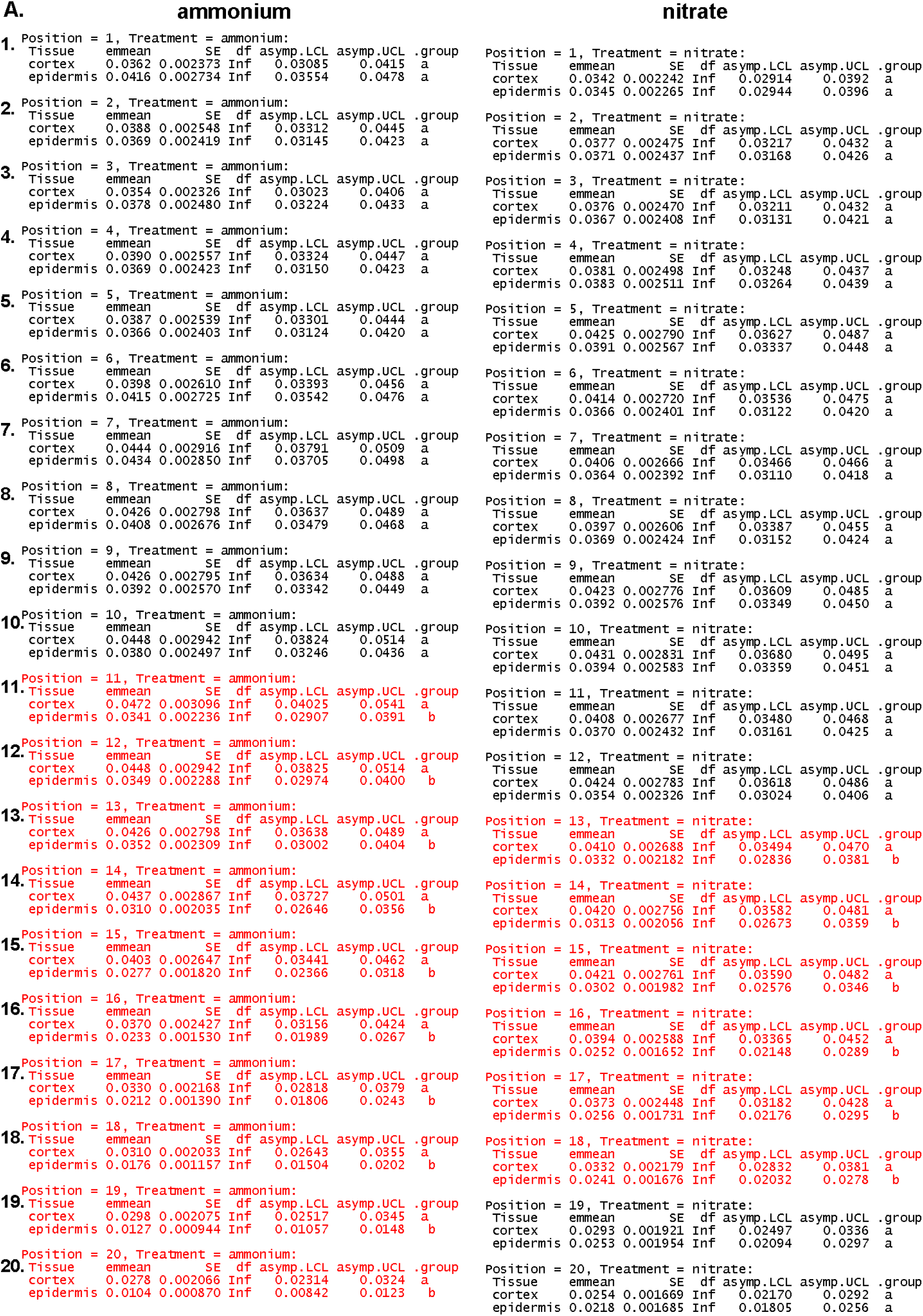

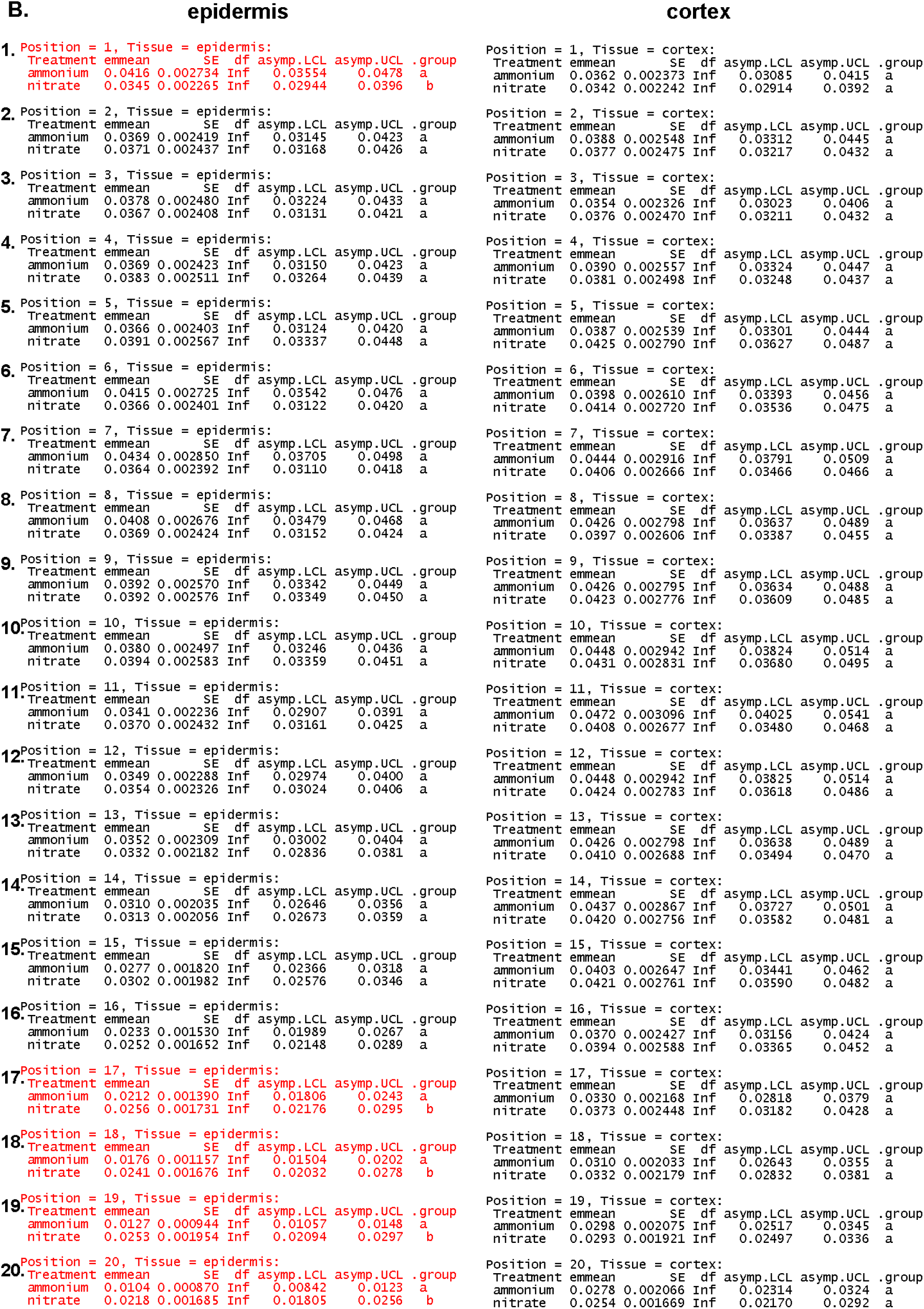

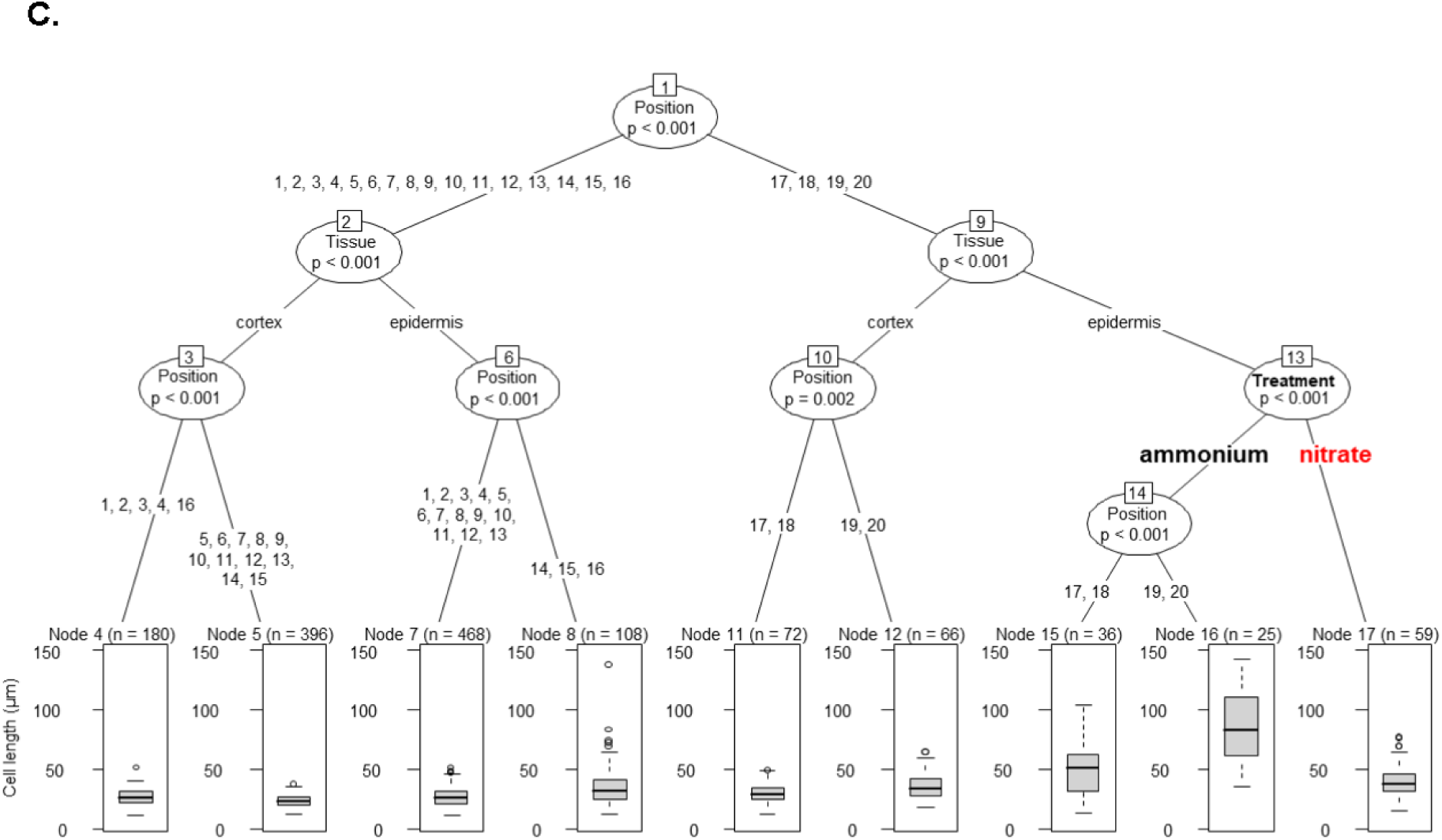

**Supplementary Document 1. Statistical reasoning**.

**A**. Estimated marginal mean (EMM) comparisons of cell lengths in different tissues (epidermis vs cortex) at each cell position (1-20 from QC) for each treatment (ammonium vs nitrate) applied on a generalized linear model (GLM). Significant differences (p<0.01) are highlighted in red. **B**. EMM comparisons of cell lengths in different treatments (ammonium vs nitrate) at each cell position (1-20 from QC) for epidermis vs cortex on the same GLM model. Significant differences (p<0.01) are highlighted in red. **C**. A decision tree based on recursive partitioning analysis shows the hierarchical importance of each treatment, tissue and cell position variable on cell length differences.

## Methods

### Plant material

*Arabidopsis thaliana* (L.) Heynh plants were used in this work. The transgenic lines *W131Y*^67^, *PIN2::PIN2-GFP*^68^ in *eir1-4* background, *PIN2::PIN2S439D-GFP, PIN2::PIN2S439A-GFP* (Vega et al.,) were introduced into *eir1-4* background; *PIN2::PIN2-Dendra*^69^, *R2D2*^70^, *DII-VENUS*^70^, *mDII-VENUS*^70^, *PIN2::nls-GFP*^71^, *DR5::LUC*^72^, *DR5::RFP*^73^, *CyclinB1::GUS*^74^ and the T-DNA mutant line *eir1-4* were described previously. *DII-VENUS* and *mDII-VENUS* in *eir1-4* background lines were obtained by manual hand pollination of the individual lines.

### Growth conditions

Seeds of *A. thaliana* were surface-sterilized by 70% ethanol and sown on a modified Murashige and Skoog (MS) medium - Boric Acid 6.2 mg/L, Calcium Chloride (anhydrous) 332.2 mg/L, Cobalt Chloride (6H_2_O) 0.025 mg/L, Cupric Sulfate (5H_2_O) 0.025 mg/L, Na_2_EDTA (2H_2_O) 37.26 mg/L, Ferrous Sulfate (7H_2_O) 27.8 mg/L, Magnesium Sulfate (anhydrous)180.7 mg/L, Molybdic Acid (disodium salt 2H_2_O) 0.25 mg/L, Potassium Iodide 0.83 mg/L, Potassium Phosphate (monobasic, anhydrous) 170 mg/L, Zinc Sulfate (7H_2_O) 8.6 mg/L – which contained 0.5mM Ammonium Succinate (Santa Cruz Biotechnology) (76 mg/L) as a nitrogen source and supplemented with 0.1% sucrose and 1% agar (Type E, Sigma A4675), pH=5,8. The nitrate amended media contained 5mM Potassium Nitrate (505 mg/L) instead of 0.5mM Ammonium Succinate. Seeds were stratified at least for 3 d and grown for 4-14 d at 21 °C in a 16 h light/8 h dark cycle.

### Root growth and root length analysis

7-day-old light-grown seedlings were transferred to either ammonium or nitrate amended plates and scanned on a daily basis for 7 days on an Epson Perfection V700 flatbed scanner. Root growth (root length changes over a given period of time) and root length were measured manually using Fiji (v1.52).

### Cell elongation and cell length analysis

Cell elongation was measured after 12 hours exposure to either to ammonium or nitrate manually with the software Fiji (v1.52).

For cell length analysis, confocal microscopic images of propidium iodide-stained *PIN2::PIN2-GFP, PIN2::PIN2S439A-GFP, PIN2::PIN2S439D-GFP*, Col-0 and *eir1-4* roots were used and the length of each cell in different cell files (epidermis and cortex) was measured manually using Fiji (v1.52).

### Imaging and image analysis

5 DAG seedlings were mounted on a slice of MS medium - containing either 0.5mM ammonium or 5mM nitrate - placed into a chambered coverslip (Lab-Tek) and imaged with Zeiss LSM700, LSM800 or LSM880 inverted confocal microscopes equipped either with a 20×/0.8 Plan-Apochromat M27 objective or a 40× Plan-Apochromat water immersion objective. Fluorescence signals for GFP (excitation 488 nm, emission 507 nm), YFP (excitation 514 nm, emission 527 nm), PI (excitation 536 nm, emission 617 nm) and DAPI (excitation 405 nm, emission 461 nm) were detected. A LaVision 2-Photon Inverted TriM Scope II from LaVision Biotec with a FLIM X16 TCSPC detector from LaVision Biotec equipped with a Olympus UApo N340 40xW, NA 1.15 was also used. Roots were observed 12 hours after transfer to ammonium or nitrate supplemented media. Long time-lapse imaging was performed using a vertically oriented LSM700 microscope as described previously^32^.

For image quantification (R2D2, DII-Venus, mDII-Venus, PIN2-GFP fluorescence intensity measurements), maximum intensity projections of confocal pictures were used. Images were handled and analysed with Fiji (v1.52) and Adobe Photoshop (Adobe Creative Cloud).

### PIN2-DENDRA photoconversion and FRAP experiments

PIN2-DENDRA experiments were executed as previously described^71^. Briefly, photoconversion of 5 DAG seedlings expressing *PIN2-Dendra* into its red form induced by illuminating the region of interest with UV light and the depletion of the red and re-apperance of the green signals in ammonium or nitrate transferred *Arabidopsis* roots was followed over time using a vertically oriented LSM700 microscope. Observation of roots initiated 10-20 minutes after transfer and images were recorded every 20 minutes (9 stacks/root/ recording). The experiment was repeated 3 times and each experiment consisted of imaging 6 roots per condition. Image analysis was performed using Fiji (v1.52). Red and green fluorescent signal changes were measured on 10-10 individual cell membranes in the TZ over a period of 6 hours. FRAP experiments were performed as described previously^75^. Briefly, individual membranes of 5 DAG old *PIN2-GFP* expressing *Arabidopsis* roots transferred either to ammonium or nitrate were bleached using the 488nm laser of a Zeiss LSM800 confocal microscope according to its built-in bleaching protocol. Recovery of the PIN2-GFP signal at the bleached areas was followed for 10 minutes and quantification of fluorescence recovery was measured using Fiji (v1.52).

### PIN2 immunodetection and staining of nuclei

For PIN2 immunostaining, 5 DAG *Arabidopsis* roots were handled as previously described^76^. Briefly, fixation was performed using 2% PFA (in 1xMTSB) supplemented with 0.1 % TritonX-100, followed by hydrophilisation using MeOH 100 % (65°C, 10 minutes), cell wall digestion using 0.2 % Driselase and 0.15 % Macerozyme in 2 mM MES, pH 5.0 (37°C, 40 minutes), and membrane permeabilisation using 3% NP-40, 10 % DMSO in 1× MTSB (37°C, 20 minutes). Anti-PIN2 (1:100) was used as a primary antibody (37°C, 120 minutes). Alexa Fluor 488 goat anti-rabbit IgG H+L (Thermo Fischer Scientific) was used as secondary antibody (1:800) (37°C, 60 minutes). Finally, samples were mounted in VECTASHIELD® Antifade Mounting Medium with DAPI (4’,6-Diamidino-2-Phenylindole, Dihydrochloride). Images were obtained using an LSM800 microscope.

### Quantification of R2D2, DII-VENUS and mDII-VENUS fluorescence signal in *Arabidopsis* roots

*R2D2* combines *RPS5A*-driven *DII* (*DII* domain of the *INDOLE-3-ACETIC ACID28* (*IAA28, DII*) from *Arabidopsis*) fused to *n3×Venus* and *RPS5A*-driven *mDII* fused to *ntdTomato* on a single transgene^40,46^. DII-VENUS is the domain II of IAA28 fused to the VENUS fast maturing YFP and mDII-VENUS is the non-degradable form of DII-VENUS. The analysis of the fluorescence intensity of either R2D2, DII-VENUS or mDII-VENUS expressing plants grown on ammonium containing and transferred on ammonium and nitrate containing medium was performed on Maximum Intensity Projection of Z-stacks of root tips acquired with a Zeiss LSM 700 inverted laser-scanning microscope as described in^60^ with slight modifications.

To quantify the fluorescence signal in each cell per selected root tissue (epidermis and cortex) first we positioned a segmented line over the nuclei in the corresponding tissues with the ROI manager tool of the software Fiji (v1.52) (Supplemental Fig 3C). Next, we analyzed the fluorescence plot profiles of the different lines with the peak analyzer function of the software Origin (OriginLab Corporation) to find local maxima along the lines, which represented the fluorescence value of the nuclei in the tissues. In case of R2D2, auxin distribution plots were derived by reciprocal mean values of the normalized n3xVenus/ntdTomato ratio. Relative auxin level data in each cell per tissue were graphed after data interpolation using the Origin built-in algorithm for smoothing.

### 3D SIM and Polar Density Analysis of PIN2-GFP

Live *Arabidopsis* seedlings, which were incubated on either nitrate or ammonium amended medium for 6-8 hours, were mounted on to coverslips as previously described by Johnson and Vert^77^ with the coverslips additionally fixed to the slide with nail polish. Cells in the elongation zone of the root epidermis were imaged using an OMX BLAZE v4 3D SIM (Applied Precision), as described^78^. Briefly, a 60x 1.42 NA Oil Immersion objective and a 100 mW 488 laser was used to make optical sections in the Z dimension, in order to capture the totality of the lateral polar domain of the subject cell. Each Z-section image is based on 15 images generated from 3 different angles and 5 different SIM patterns and reconstructed using SOFTWORX software (Applied Precision).

A maximum projection of the Z-stack was used for analysis. Images were made binary and subjected to watershed segmentation using Fiji^79^. PIN2 spots were then detected using TrackMate^80^. The number of PIN2 spots was calculated in regions of interest (0.8 microns in width times the height of the cell) at distances sequentially further away from the polar end of the cell using a custom made Matlab script. The raw number of spots in each ROI was then normalised and plotted.

### Quantification of LUCIFERASE (LUC) activity in *Arabidopsis* roots

*DR5::LUC* expressing 7 DAG *Arabidopsis* seedlings were transferred to ammonium or nitrate containing agar plates and roots (40 roots per treatment per time point) were collected after 1 and 6 HAT and snap freezed in liquid nitrogen. Frozen root tissue was extracted in Reporter Lysis Buffer (Promega) and LUC activity was measured with the Luciferase Assay Reagent (Promega) in a multiwell plate in a Biotek SynergyH1 platereader.

### Measurements of basipetal (shootward) auxin transport in *Arabidopsis* roots

The shootward transport assay of [3H]-IAA in *Arabidopsis* roots was performed according to a previous report^81^, with a few modifications. 7 DAG Col-0 or *eir1-4* seedlings were transferred to ammonium, nitrate or MS (Murashige Skoog Basal Medium) medium with 15 seedlings as one biological replicate, and 3 replicates per treatment. The [3H]-IAA (PerkinElmer, ART-0340) droplets were prepared in MS medium with 1.25% agar and 500 mM [3H]-IAA (1.45 mL in 10 mL) and were carefully placed on the root meristem (at the very end of the roots). After incubation for 6 hours in the dark, the part of the root which was covered by the droplet was cut, the remaining root parts were collected and ground completely in liquid nitrogen and homogenized in 1 mL scintillation solution (PerkinElmer, 6013199). The samples were incubated overnight to allow the radioactivity to evenly diffuse into the whole volume of the scintillation cocktail. Finally the radioactivity was measured with a scintillation counter (Hidex 300XL), with each sample counted for 100 s, 3 times. 3 samples with only the scintillation solution were used as background controls. As an additional background control another batch of samples were prepared the same way as described above except [3H]-IAA containing droplets were placed not on the root meristem but next to the seedlings. Data shown on the figure was calculated against the background.

### GUS (β-Glucuronidase) staining

*CycB1::GUS* expression was analyzed in seedling roots 7 DAG, 12 HAT to ammonium or nitrate containing media. Seedlings were incubated for 2 hours in 37°C in staining buffer containing 1mM ferricyanide, 150 mM sodium phosphate buffer (pH 7) and 1mg/ml of X-Gluc dissolved in DMSO. Seedlings were cleared using subsequent incubation at room temperature in a series of ethanol dilutions from 60% to 10% then mounted on slides with 5%ethanol-50%glycerol mounting solution. The pattern of the GUS histochemical staining was analyzed by an Olympus BX53 microscope and Olympus DP26 digital camera, controlled by cellSense Entry software.

### RT-qPCR analysis

Total RNA was extracted from excised 7 DAG roots 1, 6 and 48 HAT to ammonium or nitrate amended plates using RNeasy® Plant Mini kit (QIAGEN) according to the manufacturer’s protocol. 1µg of RNA was used to synthesize cDNA using iScriptTM cDNA synthesis kit (Bio-Rad). The analysis was carried out on a LightCycler 480 II (SW1.5.1 Version; Roche Diagnostics) with the SYBR Green I Master kit (Roche Diagnostics) according to the manufacturer’s instructions. All PCR reactions were carried out with biological and technical triplicates. Expression levels of target genes were quantified by specific primers that were designed using Quant Prime^82^, and validated by performing primer efficiency for each primers pair. The levels of expression of each gene were first measured relative to *AT4G05320* (*UBQ10*) and then to respective mock treatment.

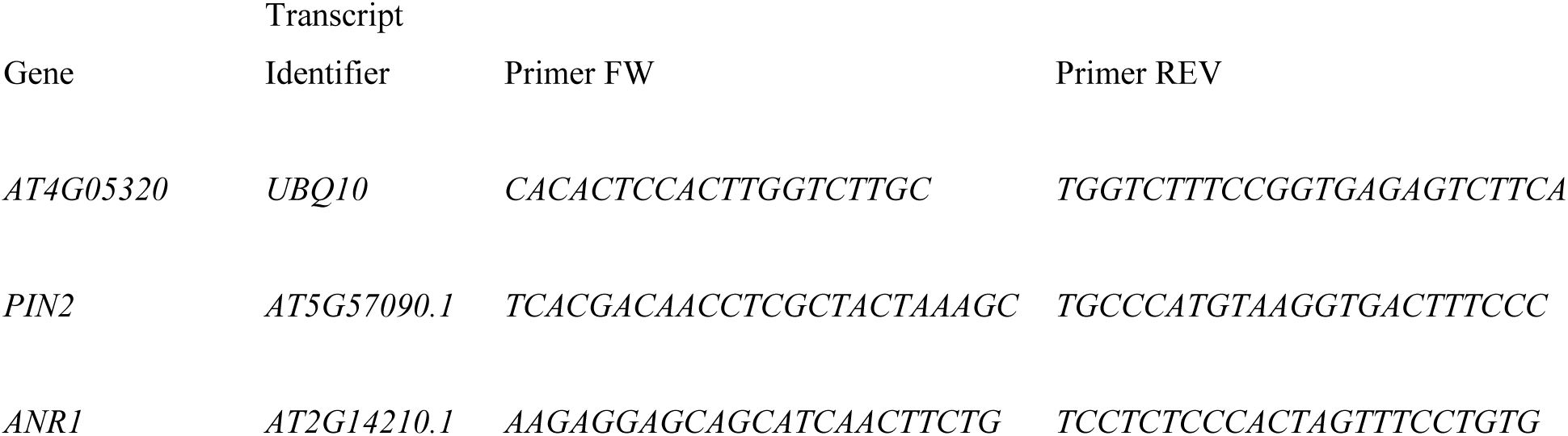

### Reproducibility and statistics

The number of independent repetitions of experiments, as well as exact sample sizes, is described in the figure legends. Statistical analysis (t-test and ANOVA) were performed using the software Origin (v2018). Statistical significance was tested as described in the figure legends.

For the regression analysis in supplementary document 1, Col-0 cell length measurements were analyzed together with associated categorical variables represented by plant sample of origin (n=18), tissue (n=2, i.e. epidermis and cortex), cell position (n=20) and treatment (n=2, i.e. NO3^-^ and NH4^+^). The importance of the variables was initially assessed via Random Forest analysis in R (v 1.2.5033). A machine learning training was conducted with the caret R package^83^ for tuning the Random Forest and the best mtry parameter was selected according to Root Mean Square Error (RMSE) and R-squared (R2) measures in R. Data distribution, skewness and kurtosis were checked with the fitdistrplus R package^84^ and Gamma distribution was chosen for setting up a regression analysis based on generalized linear models (GLMs). Besides the main effects of the variables, several models were virtually possible when the interactions between some or all the variables were considered. First, a simple model including only main effects was generated and residual vs. fitted values evaluated prior to analysis of deviance. Second, a model including main effects and all possible interactions between variables was built. The analysis of the interactions of the fit model was carried out with the phia R package^85^, showing a possible but not strong interaction between tissue and treatment factors. This insight was used to generate a third model. The performance of the second and third model was then compared by repeated k-fold cross-validation with the caret R package and the second model was selected according to RMSE and R2 measures. After analysis of deviance, post hoc pairwise comparisons were conducted with estimated marginal means (EMMs) using the emmeans R package^86^ (Supplemental Document 1a-b). A recursive partitioning analysis was performed and a decision tree was generated with the partykit R package^87^ to confirm the results showed by regression analysis and to visualize the role of different variables on cell length distribution (Supplemental Document 1c).

### Computational methods

#### Visualization of model predictions

The computer simulation representing the dynamic auxin flow through the root tissues was created using the version of VV (Vertex-Vertex) programming language and in the L-system-based modeling software L-studio^88^. The model simulates a cross-section of the plant root focusing on the cortical and epidermal tissues. Plant cells are visualized as four-sided polygons representing the cell walls. For the sake of simplicity, cell membranes and the extracellular space shared by adjacent cells are not rendered. Only the first ∼20 cells (counting from the QC) are visualized, mirroring the available experimental measurements. Meristematic and elongating cells are distinguished with different cell wall coloring; blue for meristem and yellow for elongation zone, respectively. Auxin is represented as filled green circles inside each cell, the radius of the circle proportional to the size of the cell indicates the amount of auxin present in that cell. PIN2 protein localization on the PM is represented as red dots close to the cell walls; PIN2 can be apical (shootward), basal (rootward) or lateral (outer). Despite being taken into account for mathematical calculations, cytoplasmic accumulation of PIN2 is not shown in the model visualizations. Our model enables dynamic simulation of root growth, elongation and auxin flow through the root apex. Individual cells grow, elongate and consequently divide. Auxin is pumped across cell walls through the ATP-dependent action of PIN2 proteins on the cell membrane. Auxin that reaches the outer limit of the tissues is simply removed from the system. PIN2 is expressed, trafficked and degraded according to the model rules described in the following sections.

#### Mathematical model description

The model assumes that the epidermis contributes to an active passage auxin into deeper tissues. Two main sources of auxin into the epidermis were considered:

1. The cell that is closest to the QC, which is known to be a main source of auxin production^89^.
2. The lateral root cap, which due to its structural conformation force the influx of auxin into the initial cells of the epidermis^90^.

The ordinary differential equation describing auxin dynamic in a single cell *i* is:

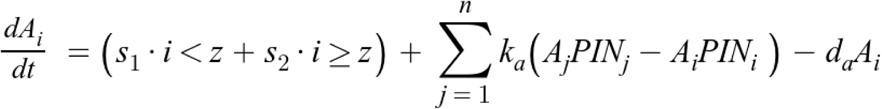

Where *s*_*1*_ and *s*_*2*_ denote the two auxin sources into the epidermis), while *z* indicates the cell location of the LRC-derived auxin influx (cell number 20 from the QC). *k*_*a*_ represents the rate of active auxin transport between cells via PIN2. The exchange of auxin occurs for each cell *j* connected to cell *i*. General processes of auxin degradation like conjugation and oxidation are summarized by a single degradation rate, *d*_*a*_.

PIN2 is the only auxin efflux carries considered in this model. High auxin concentrations lead to an increased degradation of PIN proteins^48,91^. We modeled the effect of auxin on cytoplasmic PIN2 inside cell *i* by approximating functional forms, in what follow:

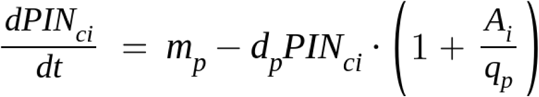

The expression parameter *m*_*p*_ indicates the basal rate of PIN2 protein synthesis. PIN2 degradation is modeled over a constant rate of degradation, *d*_*p*_, which increases linearly according to auxin levels by *q*_*p*_.

PIN2 trafficking to the apical/basal membranes is modeled as follows:

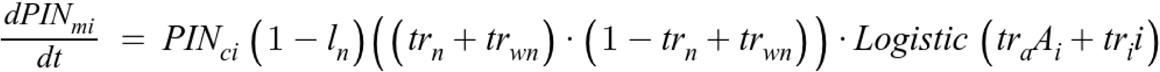

The amount of PIN2 on the membrane *m* of cell *i* is regulated by the basal trafficking rates on NO_3_^-^ (*tr*_*n)*_ or NH_4_ ^+^ (*tr*_*wn*_), which in turn is allowed to saturate to zero or to the maximum rate depending on the level of auxin and the distance from the QC, according to logistic coefficients *tr*_*a*_ and *tr*_*i*_, respectively. *l*_*n*_ represents the percentage of PIN2 that is redirected to the lateral membranes, depending on nitrate levels. In this model, nitrate level is represented as a binary variable: NO_3_^-^ for nitrate supplement and 0 for NH_4_^+^ supplement.

Cell division is regulated though a hypothetical division factor as proposed in a previous study^92^. The concentration of division factor in a single cell *i* is describes as:

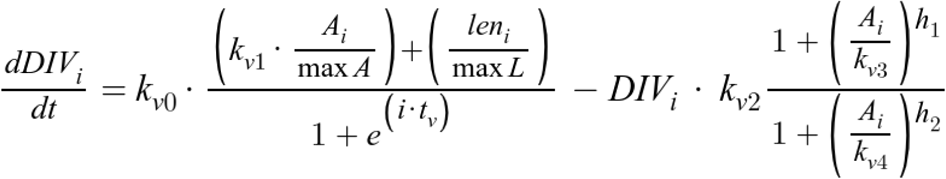

Where *k*_*v0*_ denotes the maximal synthesis rate of division factor; *len*_*i*_ and *maxL* the length of cell *i* and the maximum cell length achievable, respectively; *t*_*v*_ is the tolerance factor restricting the location in the meristem where division takes place; *k*_*v1*_ is the level of auxin-dependent division factor activation. The right part of the formula describes the hypothesized process of division factor degradation, where *k*_*v2*_ is the degradation rate of the division factor; *k*_*v3*_ and *k*_*v4*_ are the level of auxin-dependent division factor activation and saturation, respectively; *h*_*1*_ and *h*_*2*_ are hill’s coefficient.

Cell growth is an auxin-dependent mechanism and cell entrance in the elongation phase is triggered by an auxin concentration threshold. Both in the meristem and the elongation zone cell growth is defined as follow:

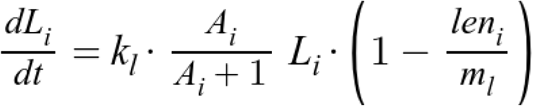

Where *k*_*l*_ indicates the cell elongation rate (depending whether the cell in the meristem or in the elongation zone), *len*_*i*_ the cell length, and *m*_*l*_ the maximum length the cell can achieve (depending whether the cell in the meristem or in the elongation zone).

#### Statistical inference and parameters estimation

Data analysis and plotting was performed using the R language environment for statistical computing^93^ and the plotting package ggplot^94^. Parameters estimation of the previously described models was carried out with the RStan^95^ and brms^96^ packages, which implement a modified version of Hamiltonian Monte Carlo sampling algorithm to approximate the parameters posterior distribution. Model comparison was performed using the loo package^97^ to carry out Pareto smoothed importance-sampling leave-one-out cross-validation (PSIS-LOO) for posterior predictive performance estimation.

#### Auxin source implementation and testing

To test auxin source impact on the model predictions we considered four possible scenarios (Supplementary Fig. 10f):

1. Model A: A naive model assuming a uniform source of auxin along the epidermis (uniform source)
2. Model B: The current model that consider two separate sources from the QC and the LRC (LRC source)
3. Model C: A highly complex model that assume input source modeled as a versatile spline (spline source)
4. Model D: A more complex but less realistic model allowing for different input of auxin for each epidermis cell (multiple point source)

To identify the best model, we tested Models A-D against experimental measurements and generate the information criteria based on the expected log probability density (Supplementary Fig. 10f). A lowest information criterion was found for Model B as indicated the posterior predictive performance, thereby Model B was used for the further study. We decided to exclude the existence of a significant influx of auxin into the cortical cells; this was backed by previous researches which suggested that at high auxin levels endodermal cells have the tendency to lateralize toward the internal tissues and not toward the cortex^98^.

#### Parameters values used in the model

Parameters values used in the model are listed below with their estimated mean and lower/upper 95% credible intervals.

| Parameter | Mean | l-95% CI | u-95% CI | Reference (when not estimated from data) |
| --- | --- | --- | --- | --- |
| $d_a / k_a$ | **0.018** | **0.018** | **0.018** | <sup>92</sup> |
| $s_1$ | **8.36** | **0.44** | **21.54** | - |
| $s_2$ | **22.53** | **9.13** | **36.46** | - |
| $z$ | **10.6** | **8.39** | **12.02** | - |
| $m_p$ | **30.49** | **16.69** | **46.00** | - |
| $d_p$ | **0.065** | **0.051** | **0.079** | - |
| $q_p$ | **100** | **100** | **100** | <sup>99</sup> |
| $l_n$ | **0.60** | **0.51** | **0.69** | - |
| $tr_n$ | **0.246** | **0.226** | **0.267** | - |
| $tr_{wn}$ | **0.13** | **0.11** | **0.15** | - |
| $tr_a$ | **-0.05** | **-0.10** | **-0.00** | - |
| $tr_i$ | **0.30** | **0.25** | **0.34** | - |
| $k_{v0}$ | **1.5** | **1.5** | **1.5** | <sup>92</sup> |
| $k_{v1}$ | **20** | **20** | **20** | <sup>92</sup> |
| $k_{v2}$ | **0.3** | **0.3** | **0.3** | <sup>92</sup> |
| $k_{v3}$ | **3.5** | **3.5** | **3.5** | <sup>92</sup> |
| $k_{v4}$ | **0.5** | **0.5** | **0.5** | <sup>92</sup> |
| $t_v$ | **0.1** | **0.1** | **0.1** | <sup>92</sup> |
| $h_1$ | **2** | **2** | **2** | <sup>92</sup> |
| $h_2$ | **3** | **3** | **3** | <sup>92</sup> |
| $k_l$ | **0.3** | **0.2** | **0.4** | <sup>100</sup> |
| $m_l$ | **200** | **150** | **250** | <sup>79</sup> |

## References

1. López-Bucio, J., Cruz-Ramírez, A. & Herrera-Estrella, L. The role of nutrient availability in regulating root architecture. Curr. Opin. Plant Biol. 6, 280–287 (2003).

2. Marhava, P. et al. Re-activation of Stem Cell Pathways for Pattern Restoration in Plant Wound Healing. Cell 177, 957-969.e13 (2019).

3. Fendrych, M., Leung, J. & Friml, J. TIR1/AFB-Aux/IAA auxin perception mediates rapid cell wall acidification and growth of Arabidopsis hypocotyls. eLife 5, (2016).

4. Marschner, H. Mineral nutrition of higher plants. (Acad. Press, 2008).

5. von Wirén, N., Gazzarrini, S. & Frommer, W. B. Regulation of mineral nitrogen uptake in plants. Plant Soil 196, 191–199 (1997).

6. Jia, Z. & Wirén, N. von. Signaling pathways underlying nitrogen-dependent changes in root system architecture: from model to crop species. J. Exp. Bot. (2020) doi:10.1093/jxb/eraa033.

7. Waidmann, S., Sarkel, E. & Kleine-Vehn, J. Same same, but different: growth responses of primary and lateral roots. J. Exp. Bot. (2020) doi:10.1093/jxb/eraa027.

8. Gruber, B. D., Giehl, R. F. H., Friedel, S. & Wirén, N. von. Plasticity of the Arabidopsis Root System under Nutrient Deficiencies. Plant Physiol. 163, 161–179 (2013).

9. Forde, B. G. Nitrogen signalling pathways shaping root system architecture: an update. Curr. Opin. Plant Biol. 21, 30–36 (2014).

10. Giehl, R. F. H. & Wirén, N. von. Root Nutrient Foraging. Plant Physiol. 166, 509–517 (2014).

11. Remans, T. et al. The Arabidopsis NRT1.1 transporter participates in the signaling pathway triggering root colonization of nitrate-rich patches. Proc. Natl. Acad. Sci. 103, 19206–19211 (2006).

12. Lima, J. E., Kojima, S., Takahashi, H. & von Wirén, N. Ammonium triggers lateral root branching in Arabidopsis in an AMMONIUM TRANSPORTER1;3-dependent manner. Plant Cell 22, 3621–3633 (2010).

13. Fredes, I., Moreno, S., Díaz, F. P. & Gutiérrez, R. A. Nitrate signaling and the control of Arabidopsis growth and development. Curr. Opin. Plant Biol. 47, 112–118 (2019).

14. Guan, P. Dancing with Hormones: A Current Perspective of Nitrate Signaling and Regulation in Arabidopsis. Front. Plant Sci. 8, 1697 (2017).

15. Ristova, D. et al. Combinatorial interaction network of transcriptomic and phenotypic responses to nitrogen and hormones in the Arabidopsis thaliana root. Sci. Signal. 9, rs13 (2016).

16. Krouk, G. Hormones and nitrate: a two-way connection. Plant Mol. Biol. 91, 599–606 (2016).

17. Chen, F. et al. Evaluation of the yield and nitrogen use efficiency of the dominant maize hybrids grown in North and Northeast China. Sci. China Life Sci. 56, 552–560 (2013).

18. Tian, Q., Chen, F., Liu, J., Zhang, F. & Mi, G. Inhibition of maize root growth by high nitrate supply is correlated with reduced IAA levels in roots. J. Plant Physiol. 165, 942–951 (2008).

19. Caba, J. M., Centeno, M. L., Fernández, B., Gresshoff, P. M. & Ligero, F. Inoculation and nitrate alter phytohormone levels in soybean roots: differences between a supernodulating mutant and the wild type. Planta 211, 98–104 (2000).

20. Tamaki, V. & Mercier, H. Cytokinins and auxin communicate nitrogen availability as long-distance signal molecules in pineapple (Ananas comosus). J. Plant Physiol. 164, 1543–1547 (2007).

21. Krouk, G. et al. Nitrate-regulated auxin transport by NRT1.1 defines a mechanism for nutrient sensing in plants. Dev. Cell 18, 927–937 (2010).

22. Ma, W. et al. Auxin biosynthetic gene TAR2 is involved in low nitrogen-mediated reprogramming of root architecture in Arabidopsis. Plant J. Cell Mol. Biol. 78, 70–79 (2014).

23. Walch-Liu, P., Liu, L.-H., Remans, T., Tester, M. & Forde, B. G. Evidence that L-glutamate can act as an exogenous signal to modulate root growth and branching in Arabidopsis thaliana. Plant Cell Physiol. 47, 1045–1057 (2006).

24. Gutiérrez, R. A. et al. Qualitative network models and genome-wide expression data define carbon/nitrogen-responsive molecular machines in Arabidopsis. Genome Biol. 8, R7 (2007).

25. Gifford, M. L., Dean, A., Gutierrez, R. A., Coruzzi, G. M. & Birnbaum, K. D. Cell-specific nitrogen responses mediate developmental plasticity. Proc. Natl. Acad. Sci. U. S. A. 105, 803–808 (2008).

26. Vidal, E. A. et al. Nitrate-responsive miR393/AFB3 regulatory module controls root system architecture in Arabidopsis thaliana. Proc. Natl. Acad. Sci. U. S. A. 107, 4477–4482 (2010).

27. Vidal, E. A., Moyano, T. C., Riveras, E., Contreras-López, O. & Gutiérrez, R. A. Systems approaches map regulatory networks downstream of the auxin receptor AFB3 in the nitrate response of Arabidopsis thaliana roots. Proc. Natl. Acad. Sci. U. S. A. 110, 12840–12845 (2013).

28. Vidal, E. A. et al. Integrated RNA-seq and sRNA-seq analysis identifies novel nitrate-responsive genes in Arabidopsis thaliana roots. BMC Genomics 14, 701 (2013).

29. Tsay, Y. F., Schroeder, J. I., Feldmann, K. A. & Crawford, N. M. The herbicide sensitivity gene CHL1 of Arabidopsis encodes a nitrate-inducible nitrate transporter. Cell 72, 705–713 (1993).

30. Mounier, E., Pervent, M., Ljung, K., Gojon, A. & Nacry, P. Auxin-mediated nitrate signalling by NRT1.1 participates in the adaptive response of Arabidopsis root architecture to the spatial heterogeneity of nitrate availability. Plant Cell Environ. 37, 162–174 (2014).

31. Gifford, M. L. et al. Plasticity Regulators Modulate Specific Root Traits in Discrete Nitrogen Environments. PLoS Genet. 9, (2013).

32. von Wangenheim, D. et al. Live tracking of moving samples in confocal microscopy for vertically grown roots. eLife 6, (2017).

33. Yazdanbakhsh, N., Sulpice, R., Graf, A., Stitt, M. & Fisahn, J. Circadian control of root elongation and C partitioning in Arabidopsis thaliana. Plant Cell Environ. 34, 877–894 (2011).

34. Baluska, F., Mancuso, S., Volkmann, D. & Barlow, P. W. Root apex transition zone: a signalling-response nexus in the root. Trends Plant Sci. 15, 402–408 (2010).

35. Kong, X., Liu, G., Liu, J. & Ding, Z. The Root Transition Zone: A Hot Spot for Signal Crosstalk. Trends Plant Sci. 23, 403–409 (2018).

36. Pavelescu, I. et al. A Sizer model for cell differentiation in Arabidopsis thaliana root growth. Mol. Syst. Biol. 14, (2018).

37. Stepanova, A. N. et al. TAA1-mediated auxin biosynthesis is essential for hormone crosstalk and plant development. Cell 133, 177–191 (2008).

38. Dello Ioio, R. et al. A genetic framework for the control of cell division and differentiation in the root meristem. Science 322, 1380–1384 (2008).

39. Blilou, I. et al. The PIN auxin efflux facilitator network controls growth and patterning in Arabidopsis roots. Nature 433, 39–44 (2005).

40. Liao, C.-Y. et al. Reporters for sensitive and quantitative measurement of auxin response. Nat. Methods 12, 207–210, 2 p following 210 (2015).

41. Goldsmith, M. H. M. The Polar Transport of Auxin. Annu. Rev. Plant Physiol. 28, 439–478 (1977).

42. Adamowski, M. & Friml, J. PIN-dependent auxin transport: action, regulation, and evolution. Plant Cell 27, 20–32 (2015).

43. Luschnig, C., Gaxiola, R. A., Grisafi, P. & Fink, G. R. EIR1, a root-specific protein involved in auxin transport, is required for gravitropism in Arabidopsis thaliana. Genes Dev. 12, 2175–2187 (1998).

44. Müller, A. et al. AtPIN2 defines a locus of Arabidopsis for root gravitropism control. EMBO J. 17, 6903–6911 (1998).

45. Hanzawa, T. et al. Cellular Auxin Homeostasis under High Temperature Is Regulated through a SORTING NEXIN1–Dependent Endosomal Trafficking Pathway[C][W]. Plant Cell 25, 3424–3433 (2013).

46. Brunoud, G. et al. A novel sensor to map auxin response and distribution at high spatio-temporal resolution. Nature (2012) doi:10.1038/nature10791.

47. Kleine-Vehn, J. et al. Recycling, clustering, and endocytosis jointly maintain PIN auxin carrier polarity at the plasma membrane. Mol. Syst. Biol. 7, 540 (2011).

48. Kleine-Vehn, J. & Friml, J. Polar Targeting and Endocytic Recycling in Auxin-Dependent Plant Development. Annu. Rev. Cell Dev. Biol. (2008) doi:10.1146/annurev.cellbio.24.110707.175254.

49. Langowski, L. et al. Cellular mechanisms for cargo delivery and polarity maintenance at different polar domains in plant cells. Cell Discov. 2, 16018 (2016).

50. Barbosa, I. C. R., Hammes, U. Z. & Schwechheimer, C. Activation and Polarity Control of PIN-FORMED Auxin Transporters by Phosphorylation. Trends Plant Sci. 23, 523–538 (2018).

51. Abas, L. et al. Intracellular trafficking and proteolysis of the Arabidopsis auxin-efflux facilitator PIN2 are involved in root gravitropism. Nat. Cell Biol. 8, 249–256 (2006).

52. Baster, P. et al. SCFTIR1/AFB-auxin signalling regulates PIN vacuolar trafficking and auxin fluxes during root gravitropism. EMBO J. 32, 260–274 (2013).

53. Tian, H., Niu, T., Yu, Q., Quan, T. & Ding, Z. Auxin gradient is crucial for the maintenance of root distal stem cell identity in Arabidopsis. Plant Signal. Behav. 8, (2013).

54. Petersson, S. V. et al. An Auxin Gradient and Maximum in the Arabidopsis Root Apex Shown by High-Resolution Cell-Specific Analysis of IAA Distribution and Synthesis. Plant Cell 21, 1659–1668 (2009).

55. Velasquez, S. M., Barbez, E., Kleine-Vehn, J. & Estevez, J. M. Auxin and cellular elongation. Plant Physiol. (2016) doi:10.1104/pp.15.01863.

56. Chapman, E. J. & Estelle, M. Cytokinin and auxin intersection in root meristems. Genome Biol. (2009) doi:10.1186/gb-2009-10-2-210.

57. Kong, X., Liu, G., Liu, J. & Ding, Z. The Root Transition Zone: A Hot Spot for Signal Crosstalk. Trends in Plant Science (2018) doi:10.1016/j.tplants.2018.02.004.

58. Jackson, R. B. & Caldwell, M. M. The Scale of Nutrient Heterogeneity Around Individual Plants and Its Quantification with Geostatistics. Ecology 74, 612–614 (1993).

59. Nacry, P., Bouguyon, E. & Gojon, A. Nitrogen acquisition by roots: physiological and developmental mechanisms ensuring plant adaptation to a fluctuating resource. Plant Soil 370, 1–29 (2013).

60. Di Mambro, R. et al. Auxin minimum triggers the developmental switch from cell division to cell differentiation in the Arabidopsis root. Proc. Natl. Acad. Sci. U. S. A. 114, E7641–E7649 (2017).

61. Barbez, E., Dünser, K., Gaidora, A., Lendl, T. & Busch, W. Auxin steers root cell expansion via apoplastic pH regulation in Arabidopsis thaliana. Proc. Natl. Acad. Sci. 114, E4884–E4893 (2017).

62. PNAS Plus: Auxin steers root cell expansion via apoplastic pH regulation in Arabidopsis thaliana. https://www.ncbi.nlm.nih.gov/pmc/articles/PMC5474774/.

63. Su, S.-H., Gibbs, N. M., Jancewicz, A. L. & Masson, P. H. Molecular Mechanisms of Root Gravitropism. Curr. Biol. CB 27, R964–R972 (2017).

64. Zwiewka, M., Bilanovičová, V., Seifu, Y. W. & Nodzyński, T. The Nuts and Bolts of PIN Auxin Efflux Carriers. Front. Plant Sci. 10, (2019).

65. Zhang, H. et al. Quantitative phosphoproteomics after auxin-stimulated lateral root induction identifies an SNX1 protein phosphorylation site required for growth. Mol. Cell. Proteomics MCP 12, 1158–1169 (2013).

66. Kumar, S., Stecher, G., Li, M., Knyaz, C. & Tamura, K. MEGA X: Molecular Evolutionary Genetics Analysis across Computing Platforms. Mol. Biol. Evol. 35, 1547–1549 (2018).

67. Geldner, N. et al. Rapid, combinatorial analysis of membrane compartments in intact plants with a multicolor marker set. Plant J. Cell Mol. Biol. 59, 169–178 (2009).

68. Vieten, A. et al. Functional redundancy of PIN proteins is accompanied by auxin-dependent cross-regulation of PIN expression. Dev. Camb. Engl. 132, 4521–4531 (2005).

69. Jásik, J. et al. PIN2 Turnover in Arabidopsis Root Epidermal Cells Explored by the Photoconvertible Protein Dendra2. PLOS ONE 8, e61403 (2013).

70. Brunoud, G. et al. A novel sensor to map auxin response and distribution at high spatio-temporal resolution. Nature 482, 103–106 (2012).

71. Salanenka, Y. et al. Gibberellin DELLA signaling targets the retromer complex to redirect protein trafficking to the plasma membrane. Proc. Natl. Acad. Sci. U. S. A. 115, 3716–3721 (2018).

72. Ulmasov, T., Murfett, J., Hagen, G. & Guilfoyle, T. J. Aux/IAA proteins repress expression of reporter genes containing natural and highly active synthetic auxin response elements. Plant Cell 9, 1963–1971 (1997).

73. Marin, E. et al. miR390, Arabidopsis TAS3 tasiRNAs, and Their AUXIN RESPONSE FACTOR Targets Define an Autoregulatory Network Quantitatively Regulating Lateral Root Growth. Plant Cell 22, 1104–1117 (2010).

74. Colón-Carmona, A., You, R., Haimovitch-Gal, T. & Doerner, P. Technical advance: spatio-temporal analysis of mitotic activity with a labile cyclin-GUS fusion protein. Plant J. Cell Mol. Biol. 20, 503–508 (1999).

75. Glanc, M., Fendrych, M. & Friml, J. Mechanistic framework for cell-intrinsic re-establishment of PIN2 polarity after cell division. Nat. Plants 4, 1082–1088 (2018).

76. Pasternak, T. et al. Protocol: an improved and universal procedure for whole-mount immunolocalization in plants. Plant Methods 11, 50 (2015).

77. Johnson, A. & Vert, G. Single Event Resolution of Plant Plasma Membrane Protein Endocytosis by TIRF Microscopy. Front. Plant Sci. 8, 612 (2017).

78. Hille, S., Akhmanova, M., Glanc, M., Johnson, A. & Friml, J. Relative Contribution of PIN-Containing Secretory Vesicles and Plasma Membrane PINs to the Directed Auxin Transport: Theoretical Estimation. Int. J. Mol. Sci. 19, (2018).

79. Schindelin, J. et al. Fiji: an open-source platform for biological-image analysis. Nat. Methods 9, 676–682 (2012).

80. Tinevez, J.-Y. et al. TrackMate: An open and extensible platform for single-particle tracking. Methods 115, 80–90 (2017).

81. Lewis, D. R. & Muday, G. K. Measurement of auxin transport in Arabidopsis thaliana. Nat. Protoc. 4, 437–451 (2009).

82. Arvidsson, S., Kwasniewski, M., Riaño-Pachón, D. M. & Mueller-Roeber, B. QuantPrime –a flexible tool for reliable high-throughput primer design for quantitative PCR. BMC Bioinformatics 9, 465 (2008).

83. Kuhn, M. et al. caret: Classification and Regression Training. (2020).

84. Delignette-Muller, M. L. & Dutang, C. fitdistrplus: An R Package for Fitting Distributions. J. Stat. Softw. 64, 1–34 (2015).

85. Rosario-Martinez, H. D., Fox, J. & R Core Team. phia: Post-Hoc Interaction Analysis. (2015).

86. Lenth, R., Singmann, H., Love, J., Buerkner, P. & Herve, M. emmeans: Estimated Marginal Means, aka Least-Squares Means. (2020).

87. Hothorn, T. & Zeileis, A. partykit: A Modular Toolkit for Recursive Partytioning in R. J. Mach. Learn. Res. 16, 3905–3909 (2015).

88. Karwowski, R. & Prusinkiewicz, P. The L−system−based plant−modeling environment L−studio 4.0 The L-system-based plant-modeling environment L-studio 4.0. in Proceedings of the 4th International Workshop on Functional−Structural Plant Models, (2004).

89. Stepanova, A. N. et al. TAA1-Mediated Auxin Biosynthesis Is Essential for Hormone Crosstalk and Plant Development. Cell (2008) doi:10.1016/j.cell.2008.01.047.

90. Xuan, W. et al. Cyclic programmed cell death stimulates hormone signaling and root development in Arabidopsis. Science (2016) doi:10.1126/science.aad2776.

91. Kleine-Vehn, J. et al. Differential degradation of PIN2 auxin efflux carrier by retromer-dependent vacuolar targeting. Proc. Natl. Acad. Sci. U. S. A. (2008) doi:10.1073/pnas.0808073105.

92. Mironova, V. V. et al. A plausible mechanism for auxin patterning along the developing root. BMC Syst. Biol. (2010) doi:10.1186/1752-0509-4-98.

93. R Development Core Team, R. R: A Language and Environment for Statistical Computing. R Foundation for Statistical Computing (2011). doi:10.1007/978-3-540-74686-7.

94. Ginestet, C. ggplot2: Elegant Graphics for Data Analysis. J. R. Stat. Soc. Ser. A Stat. Soc. (2011) doi:10.1111/j.1467-985x.2010.00676_9.x.

95. Carpenter, B. et al. Stan: A probabilistic programming language. J. Stat. Softw. (2017) doi:10.18637/jss.v076.i01.

96. Bürkner, P. C. brms: An R package for Bayesian multilevel models using Stan. J. Stat. Softw. (2017) doi:10.18637/jss.v080.i01.

97. Vehtari, A., Gelman, A. & Gabry, J. Practical Bayesian model evaluation using leave-one-out cross-validation and WAIC. Stat. Comput. (2017) doi:10.1007/s11222-016-9696-4.

98. Sauer, M. et al. Canalization of auxin flow by Aux/IAA-ARF-dependent feedback regulation of PIN polarity. Genes Dev. (2006) doi:10.1101/gad.390806.

99. Wabnik, K. et al. Emergence of tissue polarization from synergy of intracellular and extracellular auxin signaling. Mol. Syst. Biol. (2010) doi:10.1038/msb.2010.103.

100. Yang, X., Dong, G., Palaniappan, K., Mi, G. & Baskin, T. I. Temperature-compensated cell production rate and elongation zone length in the root of Arabidopsis thaliana. Plant Cell Environ. (2017) doi:10.1111/pce.12855.

